# ATP-dependent citrate lyase Drives Left Ventricular Dysfunction by Metabolic Remodeling of the Heart

**DOI:** 10.1101/2024.06.21.600099

**Authors:** Shijie Liu, Seth T. Gammon, Lin Tan, Yaqi Gao, Kyoungmin Kim, Ian K. Williamson, Janet Pham, Angela Davidian, Radhika Khanna, Benjamin D. Gould, Rebecca Salazar, Heidi Vitrac, An Dinh, Evan C. Lien, Francisca N. de L. Vitorino, Joanna M. Gongora, Sara A. Martinez, Czer S. C. Lawrence, Evan P. Kransdorf, David Leffer, Blake Hanson, Benjamin A. Garcia, Matthew G. Vander Heiden, Philip L. Lorenzi, Heinrich Taegtmeyer, David Piwnica-Worms, James F. Martin, Anja Karlstaedt

**Affiliations:** Department of Molecular Physiology and Biophysics, Baylor College of Medicine, Texas Heart Institute, Houston, TX 77030, USA; Department of Cancer Systems Imaging, The University of Texas MD Anderson Cancer Center, Houston, TX 77030, USA; Department of Bioinformatics and Computational Biology, The University of Texas MD Anderson Cancer Center, Houston, TX 77054, USA; Department of Cardiology, The Smidt Heart Institute, Cedars-Sinai Medical Center, Los Angeles, CA 90048, USA; Department of Internal Medicine, Division of Cardiology, McGovern Medical School, The University of Texas Health Science Center at Houston, Houston, TX 77030, USA; Bruker Daltonics, Billerica, MA 01821, USA; Center for Infectious Diseases, University of Texas Health Science Center, School of Public Health, Houston, TX 77030, USA; Koch Institute for Integrative Cancer Research, Massachusetts Institute of Technology, Cambridge, MA 02139-4307, USA; Department of Biochemistry and Molecular Biophysics, Washington University School of Medicine, St. Louis, MO 63110, USA; Department of Cardiothoracic Surgery, The Smidt Heart Institute, Cedars-Sinai Medical Center, Los Angeles, CA 90048, USA; Department of Epidemiology, Human Genetics, and Environmental Sciences, The University of Texas Health Science Center at Houston, Houston, TX 77030, USA; Department of Metabolism and Nutritional Programming, Van Andel Institute, Grand Rapids, MI 49503, USA; UC Department of Pediatrics, Cincinnati Children’s Hospital Medical Center, Cincinnati, OH 45229, USA; Department of Medicine, Cleveland Clinic Florida, Weston, FL 33331, USA; Dana-Farber Cancer Institute, Boston, MA 02115, USA

**Keywords:** ATP-dependent citrate lyase, Cardio-oncology, Metabolism, Systems Biology

## Abstract

**Background:** Metabolic remodeling is a hallmark of the failing heart. Oncometabolic stress during cancer increases the activity and abundance of the ATP-dependent citrate lyase (ACL, *Acly*), which promotes histone acetylation and cardiac adaptation. ACL is critical for the de novo synthesis of lipids, but how these metabolic alterations contribute to cardiac structural and functional changes remains unclear.

**Methods:** We utilized human heart tissue samples from healthy donor hearts and patients with hypertrophic cardiomyopathy. Further, we used CRISPR/Cas9 gene editing to inactivate *Acly* in cardiomyocytes of MyH6-Cas9 mice. *In vivo,* positron emission tomography and *ex vivo* stable isotope tracer labeling were used to quantify metabolic flux changes in response to the loss of ACL. We conducted a multi-omics analysis using RNA-sequencing and mass spectrometry-based metabolomics and proteomics. Experimental data were integrated into computational modeling using the metabolic network CardioNet to identify significantly dysregulated metabolic processes at a systems level.

**Results:** Here, we show that in mice, ACL drives metabolic adaptation in the heart to sustain contractile function, histone acetylation, and lipid modulation. Notably, we show that loss of ACL increases glucose oxidation while maintaining fatty acid oxidation. *Ex vivo* isotope tracing experiments revealed a reduced efflux of glucose-derived citrate from the mitochondria into the cytosol, confirming that citrate is required for reductive metabolism in the heart. We demonstrate that YAP inactivation facilitates ACL deficiency. Computational flux analysis and integrative multi-omics analysis indicate that loss of ACL induces alternative isocitrate dehydrogenase 1 flux to compensate.

**Conclusions:** This study mechanistically delineates how cardiac metabolism compensates for suppressed citrate metabolism in response to ACL loss and uncovers metabolic vulnerabilities in the heart.

## INTRODUCTION

Metabolic adaptation in the failing heart is critical to sustaining energy provision for cardiac contractile function.^1^ Increased glucose uptake and oxidation during heart failure development have been acknowledged for decades as a sign of impaired mitochondrial metabolism. Understanding the metabolic phenotype of the failing heart is incomplete and based mainly on pathogenic variants of proteins and pressure-overload models.^2–4^ Recent studies suggest that mitochondrial function and glucose metabolism support biosynthetic demands in the heart.^5, 6^ Whether cardiac metabolism requires glucose to support reductive metabolism remains unknown.

In the current understanding of cardiac metabolism, mitochondrial oxidation of glucose exclusively provides ATP for contractile function, and glycolysis replenishes glycogen stores and supports ATP provision when oxidative metabolism is limited. Studies demonstrating an increased contribution of glycolytic intermediates (e.g., glucose-6 phosphate) to support NAPDH synthesis and one-carbon metabolism have challenged these concepts, suggesting that glucose metabolism is essential in supporting reductive metabolic processes in the heart. Our previous work showed that cancer cells impair cardiac contractile function via the secretion of the oncometabolite D-2-hydroxyglutarate.^7^ One of the primary metabolic changes observed in the heart is the redirection of Krebs cycle intermediates into citrate.^7, 8^ This redirection is associated with increased activity and abundance of the enzyme ATP-dependent citrate lyase (ACL, *Acly*), which catalyzes the conversion of citrate to acetyl-CoA and oxaloacetate. ACL is critical for endogenous lipid synthesis and requires ATP, linking its function directly to cellular energy provision. Due to its involvement in cellular acetyl-CoA synthesis, ACL has been related to the regulation of histone acetylation in cancer cells^9–11^ and adipose tissue.^9, 12–15^ Changes in acetyl-CoA availability correlate with the acetylation of histone 3 at lysine residue 9 (H3K9). At the same time, oncometabolic stress is associated with increased pan-acetylation of histone 3.^7^ However, the role of ACL in the heart during physiologic and pathophysiologic remodeling remains poorly defined.

A challenge for studying cardiac metabolism is the requirement for in vivo models and the need to quantify metabolic fluxes and recapitulate human phenotypes. Clustered, regularly interspaced short palindromic repeats (CRISPR)-associated (Cas)9 genomic editing has been successfully used to generate cardiac-specific transgenic mice, which express high levels of Cas9 in the heart without inferring defects.^16, 17^ These models have been applied to disease modeling of mutations in structural proteins or cardiac regeneration. However, models studying the role of metabolic enzymes in the heart are limited. Fatty acids are the heart’s primary substrate, and the fatty acid oxidation rate corresponds with cardiac workload and contractile stimulation. Previous studies demonstrated that acetyl- and malonyl-CoA levels correspond with the total fatty acid oxidation rate through the opposing actions of acetyl-CoA carboxylase and malonyl-CoA decarboxylase.^18, 19^ These studies showed a critical role for fatty acid oxidation in driving the mitochondrial metabolism of acetyl- and malonyl-CoA in the heart. The role of ACL in cardiac oxidative metabolism is not well understood.

Here, we demonstrate that loss of ACL causes left ventricular dysfunction and promotes cardiomyopathy. Moreover, knocking down *Acly* in cardiomyocytes using in vivo CRISPR/Cas9 gene editing increases glycolysis in the presence of fatty acid oxidation. Disruption of ACL promotes metabolic adaptation in the heart to sustain cardiac contractile function and energy provision. This metabolic adaptation is characterized by increased glucose uptake and oxidation, increased pentose phosphate pathway flux, increased lipid remodeling, and modulation of histone 3 acetylation. Activation of these compensatory metabolic changes is associated with YAP inactivation and an increased supply of exogenous fatty acids to maintain the synthesis of cytosolic acetyl-CoA, as indicated by computational modeling and tracer-based flux analysis. Our multi-omics analysis revealed an unexpected finding: ACL activity is critical for cardiac contractile function and modulates lipid synthesis in the heart.

## METHODS

### Data availability

The sequencing data in this article have been deposited in the Gene Expression Omnibus under accession number GSE####[*Note: will be added after acceptance of manuscript*].

Metabolomics raw data files are available in the Supplemental Data Spreadsheet.

Visualization of metabolic flux distributions are available as a Cytoscape Network file (*.cys).

### Animals

Mouse studies were conducted by the institutional animal care and use committee (IACUC) at Cedars Sinai Medical Center (Los Angeles, CA, USA), The University of Texas Health Science Center (Houston, TX, USA), MD Anderson Cancer Center (Houston, TX, USA) and at Baylor College of Medicine (Houston, TX, USA). Myh6-Cas9 mice (background strain: C57BL/6) were a gift from Dr. Eric Olson’s lab and have been described previously.^16, 17^ Wildtype (WT) mice (C57BL/6) were purchased from The Jackson Laboratory (strain catalog number: 000664, RRID:IMSR JAX:000664). Both male and female mice were used in the study. All *in vivo* and *ex vivo* experiments were performed by IACUC approval at Cedars Sinai Medical Center (Los Angeles, CA, USA), The University of Texas Health Science Center (Houston, TX, USA), MD Anderson Cancer Center (Houston, TX, USA) or at Baylor College of Medicine (Houston, TX, USA). Mice had free access to water and food, which consisted of standard chow. Animals were housed in colony cages (maximum of 5 animals per cage) with 12-hour shifts of the light-dark cycle. The genetic Background of the mice was tested using the mini MUGA array from toe or tail biopsies through an automated genotyping provider (Transnetyx, Cordova, TN, USA).^20^ Data management and productivity planning of the mice were facilitated by Transnetyx Colony Management and Artificial Mouse Intelligence (AMI) software tools. The software tracks mouse colony breeding productivity and mouse-specific data in one place. AMI was used to develop accurate and efficient breeding plans based on breeding and research goals, assisting with the optimized production of viable mice. The software tracks breeding productivity and cage census, providing system alerts for guidance toward more efficient matings and cage footprint. Tagging was conducted at weaning using RapID tags (RapID Lab, Inc., San Francisco, CA).

### Clinical samples

All subjects were enrolled in the Smidt Heart Institute Tissue Repository (CS-IRB Pro00010979 and Pro00011910) approved by the Institutional Review Board (IRB) at Cedars-Sinai Medical Center. Informed consent was obtained from all patients. Human heart tissue samples were obtained from the patient during surgery, following the standard procedure regulated by the Smidt Heart Institute Tissue Repository.

### Quantitative real-time-PCR Analysis

Total RNA was extracted from freeze-clamped tissue (approximately 10-30 mg) using the RNeasy Plus Mini Kit for fibrous tissue (Cat#74034; Qiagen, Hilden, Germany) according to the manufacturer’s instructions. RNA concentration was determined using a Qubit 4 Fluorometer (Cat#Q33226, Invitrogen, USA; RRID:SCR_018095). cDNA was synthesized using the iScript™ cDNA Synthesis Kit according to the manufacturer’s instructions (Cat#1708890, Bio-Rad Laboratories, Hercules, CA). Quantitative real-time PCR was conducted using SYBR Green probes and measured using the QuantStudio using the following PCR settings: stage 1, 50°C 2 min, 95°C 10 min; stage 2, 95°C 15 sec, 60°C 1 min, cycles 40. Results were analyzed as ΔΔCt and expressed as the fold change in transcript levels. Reference gene expression was determined using the BioRAD reference gene panel.

### Adeno-Associated Virus (AAV) 9 Production and Delivery

Two guide RNAs (gRNA) against mouse Acly were designed using Benchling (https://benchling.com). gRNA activity was first tested in P19 cells using the PX-458 backbone, which contains a gRNA scaffold and SpCas9 (Plasmid #48183, Addgene, Watertown, MA). A surveyor assay was used for testing the activity of gRNA according to the manufacturer’s instructions (Cat#1075931, Integrated DNA Technologies, Coralville, IA). The first gRNA was cloned into the 1179_pAAV-U6-BbsI-gRNA-CB-EmGFP backbone (Plasmid#89060, Addgene, Watertown, MA; RRID:Addgene_89060) using BbsI cloning sites. The second gRNA, including the U6 promoter and gRNA scaffold, was inserted behind the first gRNA scaffold to generate a plasmid with two gRNAs for Acly. The plasmid was then submitted to the IDDRC Neuroconnectivity Core at Baylor College of Medicine (Houston, TX) for virus packaging. MyH6-Cas9 mice were subcutaneously injected with 1 x10^11^ viral particles at P6. Mice were then subjected to echocardiography at 12, 14, 16, 18, and 20 weeks. The Acly gRNA sequences were as follows: (1) Acly-RNA1: GAGAGAGATTGACCCCGACG and (2) Acly-RNA2: TCCTGGCTAAAACCTCGCCT.

### Transthoracic Echocardiography

Cardiac function was determined using transthoracic echocardiography (Vevo2100, SW Version 2.2.0; SW Build 12088, 40 MHz, 550S probe, FUJIFILM VisualSonics, Toronto, Canada). After alignment in the transverse B-mode with the papillary muscles, cardiac function was measured on M-mode images. At least five mice were used for each group. The figure legend indicates that group sizes for in vivo studies vary. Samples sizes were adequately powered to observe the effects. Echocardiography results were analyzed blindly.

### Positron Emission Tomography/Computer tomography (PET/CT)

Mice were injected and imaged in the afternoon after 4 hours of fasting. Water was always provided ad libitum. Mice were injected with [¹⁸F]-Fluorodeoxyglucose ([¹⁸F]-FDG, 1.7 +/- 0.1 MBq, 46 µCi +/- 2.8 µCi, standard deviation). Individual injected doses, weights, and blood glucose were recorded for each mouse to correct for injected activity and individual changes in glucose potentially. Mice were scanned at the indicated times after injection with a 10-min PET/CT acquisition (Albira SI, Bruker) using a 15-cm field of view (FOV); CT images were acquired for fusion using a 7-cm FOV, stepped and stitched if required. PET reconstruction was by MLEM 3D, 12 iterations, with decay, random and scatter correction applied to 500-μm isotropic voxels. CT is reconstructed with filtered back projection and 500-μm isotropic voxels. The preclinical PET system at the MD Anderson Small Animal Imaging Facility includes daily quality control with flood phantoms and at least one annual preventive maintenance, including quantitative calibration. The actual injected dose was calculated based on measuring the pre- and post-injection activity in the syringe with a dose calibrator (Capintec), and mice were individually weighed for static images. Image data were decay-corrected to injection time (Albira SI, Bruker) and corrected for individual injected dose (%ID/cc) and, where indicated, normalized to animal weight, expressed as SUV (g/cc) (PMOD, PMOD Technologies).

### Protein Analysis

The expression of proteins was analyzed using Western blotting. Proteins were extracted from flash-frozen tissues as described previously.^7, 21^ Briefly, tissue samples were homogenized and lysed in RIPA buffer (10 mmol/L Tris at pH7.5, 1 mmol/L EDTA, 0.1% SDS, 1% Triton X-100, 0.1% Sodium deoxycholate, 5 mol/L NaCl) containing EDTA-free Protease and Phosphatase inhibitors (Cat#11873580001, Rose Holding AG, Basel, Switzerland). Proteins were separated on 4-12% SDS-PAGE gels (Precast Protein gels, Cat#NP0322BOX, Invitrogen, Waltham, MA), transferred to PVDF membranes and probed with antibodies from Cell Signaling Technology (CS, Danvers, MA, USA) against ACL (Cat#4332, Cell Signaling Technology, Danvers, MA; RRID:AB_2223744), total YAP/TAZ (Cat#8418T, Cell Signaling Technology, Danvers, MA; RRID: AB_10950494), Phospho-YAP (Ser127) (Cat#13008, Cell Signaling Technology, Danvers, MA; RRID: AB_2650553), Phospho-YAP (Ser397) (Cat#13619T, Cell Signaling Technology, Danvers, MA; RRID: AB_2650554), Acyl-CoA synthetase short chain family member 1 (ACSS1) (Cat#3658T, Cell Signaling Technology, Danvers, MA;RRID:AB_2222710), Acyl-CoA synthetase short chain family member 2 (ACSS2) (Cat#ab133664, Abcam; RRID:AB_2943489). Protein levels were detected by immunoblotting using horseradish peroxidase-conjugated secondary antibodies and chemiluminescence. Secondary antibodies: goat anti-rabbit IgG (Cat#7074, Cell Signaling Technology, Danvers, MA) and goat anti-rabbit IgG (Cat#7076, Cell Signaling Technology, Danvers, MA).

### Cardiomyocyte Isolations

Adult mouse ventricular cardiomyocytes (AMVMs) were isolated as described previously ^21^. Male and female mice (20-week-old) were anesthetized using nembutal intraperitoneal injections, and their hearts were excised. The aorta was cannulated, and hearts were perfused retrogradely for 15-18 min at 37°C with HEPES-buffer containing 100 mg/mL collagenase type 2 (Cat#9001-12-1, Worthington, USA) and digested. At the end of the perfusion protocol, hearts were mechanically sheared and filtered through a 100-mm mesh filter (Cat#431752, Corning, Durham, NC). The single-cell suspension was centrifuged at 20 g for 3 minutes. Cell pellets were enriched for cardiomyocytes by washing twice with HEPES buffer containing CaCl_2_ and centrifugation at 20 g for 3 minutes.

### RNA sequencing

#### RNA extraction and library preparation

RNA-sequencing libraries were prepared using the Illumina TruSeq Stranded mRNA Sample Preparation Kit (Cat#20020594, Illumina, San Diego, CA). RNA was extracted from isolated cardiomyocytes using the QIAGEN RNeasy Fibrous Tissue Mini Kit (Cat#74704, Qiagen, Hilden, Germany). RNA concentration was determined using a Qubit 4.0 fluorometer and Qubit™ RNA HS Assay Kit (Cat#Q32852, Thermo Fisher Scientific, Waltham, MA) according to the manufacturer’s instructions. The quality of each RNA sample was checked using an Agilent 4200 Tape Station with Agilent High Sensitivity RNA ScreenTape (Cat#5067-5579, Agilent Technologies, Santa Clara, CA) according to the manufacturer’s instructions. Poly-A RNA was purified from mRNA using oligo-dT magnetic beads for library preparation. mRNA was fragmented, and cDNA was synthesized. A-tailing was conducted to enable subsequent ligation of paired-end sequencing adapters. Library quality was determined using an Agilent 4200 Tape Station with Agilent High Sensitivity D5000 DNA ScreenTape (Cat#5067-5592, Agilent Technologies, Santa Clara, CA) to ensure correct fragment sizes between 250 and 300 bp.

#### RNA sequencing and data processing

Libraries were clustered onto a HiSeq patterned flow cell using a cBot2 system (Illumina, Inc., San Diego, CA) and HiSeq 3000/4000 PE Cluster Kit (Cat#PE-410-1001, Illumina, Inc., San Diego, CA). The clustered flow cell was then run on a HiSeq 4000 Sequencing System (Illumina, Inc., San Diego, CA) using a HiSeq 3000/4000 SBS 300-cycle kit (Cat#FC-410-1003, Illumina, Inc., San Diego, CA) for 2×150 paired-end reads. All instrument preparation and run parameters were carried out or selected according to manufacturer specifications. Raw reads from the sequencer were demultiplexed and converted to fastq format using bcl2fastq v2.19.1.403 (Illumina, Inc., San Diego, CA), optioned to allow for 0 mismatches in the barcodes.

### Mouse heart perfusions

Male and female mice (20-week-old) were anesthetized using pentobarbital (100 mg/Kg, intraperitoneal), and their hearts were excised. Hearts were perfused retrogradely in the Langendorff preparation for 30 min with Krebs-Henseleit (KH) buffer containing physiologic levels of substrates (5 mM [U-^13^C]-glucose, 0.5 mM L-lactate, 1 mM glutamine). ^7, 21^ At the end of the perfusion protocol, hearts were freeze-clamped using aluminum tongs cooled in liquid nitrogen and stored at -80°C for further tissue analysis.

### Isolated-working heart perfusions

Rat hearts were perfused by the method described earlier.^22^ Briefly, rats (10-12 weeks old, 340 to 380 g) were anesthetized with pentobarbital (100 mg/kg, intraperitoneal) and heparinized (200 U) through direct injection into the inferior *V. cava* after laparotomy. Next, the chest was opened, and the heart rapidly excised and arrested in ice-cold Krebs-Henseleit (KH) buffer (120 mmol/L NaCl, 5 mmol/L KCl, 1.2 mmol/L MgSO_4_, 1.2 mmol/L KH_2_PO_4_, 25 mmol/L NaHCO_3_, 2.5 mmol/L Ca^2+^, 5 mmol/L glucose) at pH 7.4. Hearts were mounted on a cannula assembly and perfused in the working heart apparatus at 37°C with KH buffer equilibrated with 95% O_2_ – 5% CO_2_. The buffer contained glucose as the only exogenous substrate. The filling pressure was 15 cmH_2_O with an afterload of 100 cmH_2_O from the start to min 55 of the perfusion. At this time, epinephrine (1 μmol/L) was added to the buffer, and the afterload was raised to 140 cmH_2_O^22, 23^. The cardiac performance was calculated as the product of cardiac output (sum of coronary and aortic flow, m^3^/min) and the afterload (Pa). Aortic pressure and heart rate were measured continuously with a 3 French catheter (Millar Instruments, Houston, TX, USA) connected to PowerLab 8/30 recording system (ADInstruments, Colorado Springs, CO, USA). At the end of the experiments, the hearts were freeze-clamped with aluminum tongs, cooled in liquid N_2,_ and stored at -80 °C until further use.

### Histone extraction and LC-MS/MS analysis

The histones were extracted and prepared for chemical derivatization and digestion as described previously.^24, 25^ In brief, the lysine residues from histones were derivatized with the propionylation at a reagent to sample ratio 1:2 (v/v) containing acetonitrile and propionic anhydride (3:1, v/v), and the solution pH was adjusted to 8.0 using ammonium hydroxide. The propionylation was performed twice and the samples were dried on speed vac. The derivatized histones were then digested with trypsin at a 1:50 ratio (wt/wt) in 50 mM ammonium bicarbonate buffer at 37°C overnight. The N-termini of histone peptides were derivatized with the propionylation reagent twice and dried on speed vac. The peptides were desalted with the self-packed C18 stage tip. The purified peptides were then dried and reconstituted in 0.1% formic acid. An LC-MS/MS system consisted of a Vanquish Neo UHPLC coupled to an Orbitrap Exploris 240 (Thermo Scientific) was used for peptide analysis. Histones peptide samples were maintained at 7 °C on sample tray in LC. Separation of peptides was carried out on an Easy-Spray™ PepMap™ Neo nano-column (2 µm, C18, 75 µm X 150 mm) at room temperature with a mobile phase. The chromatography conditions consisted of a linear gradient from 2 to 32% solvent B (0.1% formic acid in 100% acetonitrile) in solvent A (0.1% formic acid in water) over 48 min and then 42 to 98% solvent B over 12 min at a flow rate of 300 nL/min. The mass spectrometer was programmed for data-independent acquisition (DIA). One acquisition cycle consisted of a full MS scan, 35 DIA MS/MS scans of 24 m/z isolation width starting from 295 m/z to reach 1100 m/z. Typically, full MS scans were acquired in the Orbitrap mass analyzer across 290–1100 m/z at a resolution of 60,000 in positive profile mode with an auto maximum injection time and an AGC target of 300%. MS/MS data from HCD fragmentation was collected in the Orbitrap. These scans typically used an NCE of 30, an AGC target of 1000%, and a maximum injection time of 60 ms. Histone MS data were analyzed with EpiProfile. ^26^

### Metabolite Extraction from Heart Tissue

Metabolites were isolated as described previously.^7^ Briefly, a tissue sample (5 mg) was homogenized in 500 µL ice-cold MS-grade methanol, acetonitrile, and water at a volume ratio of 40:40:20. Homogenization of cell sample was achieved by freeze-thawing sample three-times. Extracts were centrifuged at 17,000 g for 5 min at 4°C. Then the supernatant was transferred to clean glass tubes. Myristic acid D27 (Cat.No. #366889; Sigma-Aldrich, St. Louis, MO) was used as an internal standard. An aliquot of myristic acid (0.03 mg/mL) was added to each biological extract. The sample was then evaporated to dryness and stored at - 80°C for further GC-MS analysis.

### Lipid Extraction from Heart Tissue

Membrane lipids were isolated using a neutral Bligh-Dyer method. Briefly, a tissue sample (10 mg) was homogenized in 500 µL ice-cold MS-grade methanol and chloroform in a volume ratio of 66.7:33.3 to create a single-phase solution. The solution was incubated for 30 min at room temperature with intermittent mixing. After centrifugation at 3,500 x g for 10 min the supernatant was converted into a two-phase solution by chloroform. After centrifugation at 3,500 × *g* for 10 min, the lower phase was recovered and dried under a stream of nitrogen gas.

### Gas chromatography/mass spectrometry (GC/MS) analysis of polar metabolites

Metabolites were isolated as described previously.^7^ Briefly, frozen heart tissue samples were homogenized in 0.4 mL −80 °C cold MS-grade methanol, acetonitrile, and water at a volume ratio of 40:40:20 (v/v/v/). Freeze-thawing sample three times achieved further disruption of the tissue sample. Extracts were centrifuged at 13,000 rpm for 5 min at 4 °C. The supernatant was transferred to clean glass tubes, and D27-myristic acid (0.15 µg/µL, Cat. No. 366889, Sigma-Aldrich; St Louis, MO) was added as internal run standard. Vials were covered with a breathable membrane, and metabolites were evaporated to dryness under vacuum. Dried samples were stored at -80°C for further GC-MS analysis. For derivatization of polar metabolites, frozen and dried metabolites from cell extractions were dissolved in 20 µL of Methoxamine (MOX) Reagent (2% solution of methoxyamine-hydrogen chloride in pyridine; Cat. No. 89803 and 270970; Sigma-Aldrich; St Louis, MO) and incubated at 30°C for 90 minutes. Subsequently, 90 µL of N-tert-Butyldimethylsilyl-N-methyltrifluoroacetamide with 1% tert-Butyldimethyl-chlorosilane (TBDMS; Cat. No. 375934, Sigma-Aldrich; St Louis, MO) was added, and samples were incubated under shaking at 37°C for 30 minutes. GC-MS analysis was conducted using an Agilent 8890 GC coupled with an Agilent 5977-mass selective detector. Metabolites were separated on an Agilent HP-5MS Ultra Inert capillary column (Cat. No. 19091S-433UI). For each sample, 1 µL was injected at 250°C using helium gas as a carrier with a flow rate of 1.1064 mL/min. For the measurement of polar metabolites, the GC oven temperature was kept at 60°C and increased to 325°C at a rate of 10°C/min (10 min hold), followed by a post-run temperature at 325°C for 1 min. The total run time was 37 min. The MS source and quadrupole were kept at 230°C and 150°C, respectively, and the detector was run in scanning mode, recording ion abundance within 50 to 650 m/z.

### GC/MS analysis of fatty acid methyl esters

Fatty acid methyl esters (FAMEs) were analyzed by GC/MS as described by Lien et al.^27^ Dried lipid extracts were resuspended in 100 µL toluene in glass vials and derivatized with 200 µL of 2% sulfuric acid in methanol overnight at 50°C. Following derivatization, 500 μl of 5% NaCl was added, and FAMEs were extracted twice with 500 μl of hexane.

Bond Elut Florosil Solid Phase Extraction (SPE) cartridges (Agilent Technologies 12113049) were used to separate polar compounds from nonpolar matrices. After elution, FAMEs were dried under nitrogen gas, and resuspended in hexane for further analysis. GC/MS analysis was conducted with a DB-FastFAME column (Agilent Technologies G3903-63011) using an Agilent 7890A gas chromatograph coupled to an Agilent 5975C mass spectrometer. After injection, the GC oven was held at 50°C for 0.5 min, increased to 194°C at 25°C/min, held at 194°C for 1 min, increased to 245°C at 5°C/min, and held at 245°C for 3 min. The total run time was 20.5 min. The MS source and quadrupole were kept at 230°C and 150°C, respectively, and the detector was run in scanning mode, recording ion abundance within 104 to 412 m/z.

### LC-MS/MS for CoAs

Dried extracts were reconstituted in 100 µL 5mM ammonium acetate in water prior to MS, and 5 µL were injected into a Thermo Scientific TSQ Quantiva triple quadrupole mass spectrometer coupled with a Dionex UltiMate 3000 HPLC system. CoA species were separated on a Thermo Scientific Accucore C30 2.1x 150 mm column with 2.6 µm particle size (Cat. No.27826-152130). The column temperature was controlled at 35°C. Mobile phase A (MPA) was 5 mM ammonium acetate in water. Mobile phase B (MPB) was 100% methanol. The flow rate was 150 µL/min, and the gradient conditions were: initial 5% MPB, held at 5% MPB for 2 mins, then increased to 95% MPB at 6.5 min, held at 95% MPB for 3.5 min, returned to initial conditions at 10.5 min, equilibrated for 4.5 min. The total run time was 15 min. The mass spectrometer was operated in the multiple reaction monitoring (MRM) positive ion electrospray mode. The ion spray voltage was adjusted to 3600V, ion transfer tube and vaporizer temperatures were set at 350°C and 275 °C. The sheath, Auxiliary, and Sweep gas flows were at 35, 10, and 0, respectively. Raw data files were imported to TraceFinder^TM^ 5.1 SP1 (Thermo Fisher Waltham, MA, USA; Cat. No. OPTON-31001) for peak identification and quantification. Metabolite abundances were normalized by DNA concentrations. For the ^13^C tracer experiment, data were acquired using a Thermo Orbitrap Fusion Tribrid mass spectrometer coupled with a Thermo Vanquish LC system at a resolution of 240,000 with full scan mode (800-900 amu).

### Analysis of Palmitate and Palmitoylcarnitine by LC-MS

To determine the incorporation of ^13^C_16_-palmitate into palmitoylcarnitine, tissue samples from perfused mouse hearts were analyzed by ultra-high-resolution mass spectrometry (HRMS). Approximately 20-30 mg of mouse heart tissue were pulverized on liquid nitrogen, then homogenized using Precellys Tissue Homogenizer. Metabolites were extracted using 0.4 mL ice-cold methanol. Extracts were centrifuged at 17,000 g for 10 min at 4°C, supernatants were transferred to clean glass autosampler vials, and 10 μL was injected for analysis by LC-MS. LC mobile phase A (MPA) was water containing 0.1% formic acid, and mobile phase B (MPB) was acetonitrile containing 0.1% formic acid. Thermo Vanquish LC system included an Accucore C30 column (2.6 µm particle size, 150 x 2.1 mm) with a column compartment kept at 35°C. The autosampler tray was chilled to 4°C. The mobile phase flow rate was 200 µL/min, and the gradient elution program was: 0-2 min, 50% MPB; 2-10 min, 50-95% MPB; 10-15 min, held at 95% MPB; 15.0-15.5 min, 95-50% MPB; 15.5-20 min, re-equilibrate column at 50%MPB. The total run time was 20 min. Data were acquired using a Thermo Orbitrap Exploris 240 Mass Spectrometer under ESI positive and negative ionization mode (polarity switching) at a resolution 240,000 with full scan mode. Raw data files were imported to Thermo Trace Finder 5.1 software for final analysis. The fractional abundance of each isotopologue is calculated by the peak area of the corresponding isotopologue normalized by the sum of all isotopologue areas. The relative abundance of each metabolite was normalized by tissue weight.

### Flux balance analysis using CardioNet

*In silico* simulations were conducted using the metabolic network of the mammalian heart metabolism, CardioNet.^7, 19, 28^ Mathematical modelling has previously been used to study the dynamics of cardiac metabolism in response to stress, and CardioNet has been successfully applied to identify limiting metabolic processes and estimate flux distributions.^5, 7, 8, 21, 29, 30^ Flux balance analysis (FBA) allows to estimate flux rates based on metabolic constraints that are defined by the extracellular environment (e.g. oxygen and nutrient supply), cellular demands (e.g. proliferation, contraction) and tissue type. FBA uses mathematical optimization models to describe complex biological systems. Each model has the following components: parameters, variables, constraints, and objective functions. Metabolic reactions that can carry flux are defined as variables within the optimization problem. Metabolite abundances from GC/MS analysis and enzymatic activities were used as constraints for network reactions. The objective function is to maximize ATP hydrolysis as a reflection of cardiac contraction (v_ATPase_) that satisfy the metabolic constraints. Furthermore, we demanded the synthesis of biomass as defined previously.^5, 7, 21, 29^ Boundary conditions for plasma metabolites were based on previously reported values. ^5, 7, 21, 29^ We determined flux distributions (*v*_*m*_) and estimated flux rate changes (v_FC_) as described previously.^5, 19, 21, 30^ The following flux balance analysis was applied to identify steady-state flux distributions that agree with applied substrate uptake and release rates, and changes in metabolite pools:

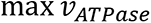

subject to

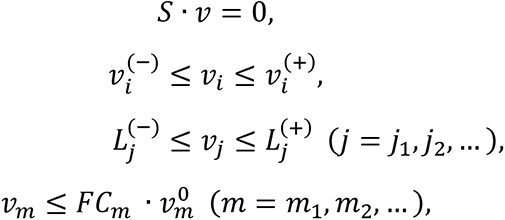

where *v*_*i*_ denotes the flux rate change through reaction *i*, *v*_*j*_ denotes the measured uptake or secretion rate through reaction *j*, *S* is the stoichiometric matrix, and 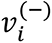 and 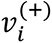 are flux constraints. The GUROBI LP solver was used to find the solution to the FBA problems.^31^

## RESULTS

### Cardiac-specific Acly deletion induces left ventricular remodeling

ACL is a critical enzyme in de novo synthesis of acetyl-CoA and regulating cytosolic citrate levels. We observed a significant reduction in cardiac ACL expression in explanted hearts from patients undergoing transplantation due to hypertrophic cardiomyopathy (HCM) compared to human heart tissue samples from normal donor hearts (**Figure 1A**). Based on these findings, we postulated that ACL plays a role in maintaining cardiac energy homeostasis. To understand the *in vivo* role of ACL in the heart, we utilized CRISPR/Cas9 gene editing to inactivate *Acly* in cardiomyocytes of MyH6-Cas9 mice (**Figure 1B**).^17^ Two guide RNAs (gRNAs) with specificity for *Acly* were delivered via associated virus 9 (AAV9) infection to regenerative-stage MyH6-Cas9^+^ and MyH6-Cas9^-^ cardiomyocytes (**Figure S1A** and **S1B**). Cardiomyocytes isolated from MyH6-Cas9^+^ mice injected with AAV9-Acly-gRNA exhibited gene editing at the *Acly* locus (**Figure S1A**). Consistently, we detected a 60% reduction of *Acly* expression in cardiomyocytes from MyH6-Cas9^+^ mice compared to MyH6-Cas9^-^ controls (**Figure 1C** and **Figure S1B**). ACL expression remained in other internal organs, including the liver, lung, and kidneys, as well as adipose tissue and skeletal muscle (**Figure 1C, Figure S1C-S1F**). Our data show that delivering AAV9-Acly-gRNA into MyH6-Cas9^+^ mice causes immediate and sustained deletion of *Acly,* exclusively in cardiomyocytes. We refer to MyH6-Cas9^+^ and MyH6-Cas9^-^ mice injected with AAV9-Acly-gRNA as Acly-knockdown (*Acly^KD^*) and WT mice, respectively. Loss of *Acly* increased total heart weight (HW) in 20-week-old male and female hearts (normalized by tibia length, TL) (**Figure 1D**). Hearts from 20-week-old male and female *Acly^KD^* mice exhibited significant cardiomyopathy (**Figure 1E**) compared to age-matched WT mice by cardiac morphology (**Figure S1G**) and cardiac function (**Figure 1E**). *Acly* deficiency markedly impaired heart function starting at 12 weeks (Figure S1H) and progressively worsened by 20 weeks. *Acly^KD^* mice showed an increased left ventricular (LV) mass by echocardiography (**Figure 1F**), decreased LV ejection fraction (LVEF) (**Figure 1G**) and decreased fractional shortening (FS) at 20 weeks (**Figure 1G**, **Table 1)**. Our data indicate that at 20 weeks, *Acly^KD^* causes a systolic LV dysfunction and eccentric cardiac remodeling, affecting both sexes. Together, these results identify *Acly* as a critical enzyme that sustains cardiac contractile function in the adult heart.

**Figure 1.**
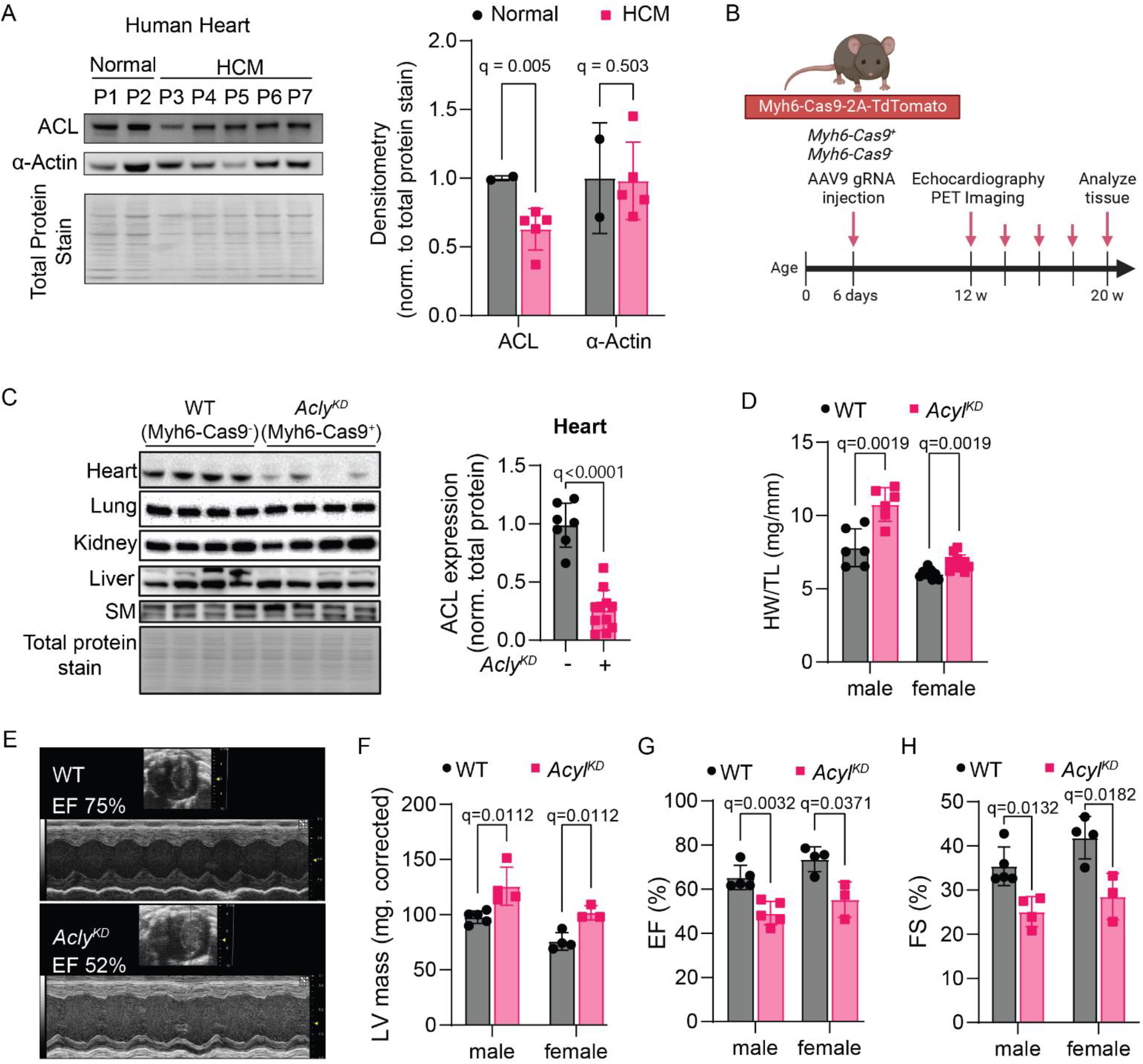
Cardiac-specific knockdown of *Acly* promotes left ventricular dysfunction. (A) ACL protein abundance in healthy human hearts (n=2; Patient (P) 1 and 2) and tissue samples from patients with hypertrophic cardiomyopathy (n=5; P3-P7). Statistical significance was calculated by unpaired t-test followed by multiple comparisons analysis using a false discovery rate < 5% by the two-step method of Benjamini, Krieger, and Yekutieli. (B) Experimental design for CRISPR/Cas9-mediated cardiac-specific knockdown (KD) of *Acly* using Myh6-Cas9-2d-tomato transgenic mice. Single guide RNAs (sgRNA) were delivered using AAV9 at P6 to achieve cardiac-specific deletion of *Acly.* Myh6-Cas9-mice retained *Acly* expression (WT, Myh6-Cas9^-^), while Myh6-Cas9^+^ lost *Acly* expression (*Acly^KD^)*. (C) Comparison of ACL protein abundance in adult mouse ventricular cardiomyocytes (AMVMs), internal organs (liver, kidney, lung), and Mus.gastrocnemius (SM) from WT (MyH6-Cas9^-^) and *Acly^KD^* mice at 20 weeks. Representative western blot images. Ponceau S stain for total protein extracts used for western blot. AMVMs: n=7-10 male and female mice/group. Statistical significance was calculated by unpaired t-test followed by multiple comparisons analysis using a false discovery rate < 5% by the two-step method of Benjamini, Krieger, and Yekutieli. (D) Heart weight (mg) normalized to tibia length (mm) from WT (MyH6-Cas9^-^) and *Acly^KD^* mice at 20 weeks. n=10-14 male and female mice/group. 2-way ANOVA. Multiple comparison analysis by Sidak. (E) Representative transthoracic echocardiography images in male and female WT (MyH6-Cas9^-^) and *Acly^KD^* mice at 20 weeks. (F) Calculated ejection fraction from male and female WT (MyH6-Cas9^-^) or *Acly^KD^* mice at 20 weeks. n = 7 male and female mice/group. Statistical significance was calculated by multiple unpaired t-tests followed by multiple comparisons analysis using a false discovery rate < 5% by the two-step method of Benjamini, Krieger, and Yekutieli. (G-H) Fractional shortening (G) and left ventricular diameters (H) indicate cardiac remodeling in male and female WT (MyH6-Cas9^-^) and *Acly^KD^* mice at 20 weeks. LVESD left ventricular end-systolic diameter; LVEDD left ventricular end-diastolic diameter. n = 7 male and female mice/group. Statistical significance was calculated by multiple unpaired t-tests followed by multiple comparisons analysis using a false discovery rate < 5% by the two-step method of Benjamini, Krieger, and Yekutieli. Numerical q-values are plotted for each comparison. Each data point represents a biological replicate, and error bars represent standard deviation.

### Loss of *Acly* drives adaption in cardiac lipid metabolism

ACL is critical for the de novo synthesis of acetyl-CoA and complex lipids in cells. It was unclear how the cardiac-specific knockdown of ACL impairs cardiac function and promotes cardiac hypertrophy. Therefore, we used transcriptomic analysis of isolated adult mouse ventricular cardiomyocytes to understand the *in vivo* adaptations in response to the loss of *Acly* in the heart. Principal component analysis (PCA) and unsupervised hierarchical clustering of normalized RNA-sequencing (RNA-seq) demonstrated a distinct separation between *Acly^KD^* and littermate WT controls (**Figure S2A** and **Figure S2B**). Gene expression analysis revealed 268 upregulated and 340 down-regulated genes (adjusted *P*<0.05) (**Figure S2C**), which are consistent with increased mitochondrial dysfunction, contractile dysfunction, and heart failure (**Figure S2D**). Gene ontology enrichment analysis of differentially regulated genes (*P*<0.01) in response to loss of *Acly* indicated transcriptomic remodeling in desaturation and elongation of fatty acids (CYB5R2, CYTB), lipid metabolic processes (PLCB2, ACOT11), mitochondrial and respiratory electron transport (ND1, ND3), and sarcomere organization (MYH7) (**Figure 2A**). Consistently predicted downregulation of transcription factors included PPARGC1A, MEF2C, and TEAD1, which are all linked to the regulation of lipid metabolism. The downregulation of genes involved in lipid synthesis and modulation was surprising, considering that de novo fatty acid synthesis is minimal in the heart. Most lipids are thought to be provided through extracellular uptake from lipoprotein particles. Therefore, we assessed the abundances of major enzymes in fatty acid synthesis and modification using RT-qPCR: (1) very long-chain fatty acid elongase 1, 5, and 7; (2) stearoyl-CoA desaturase (*Scd*) 2,3 and 4; (3) fatty acid desaturase 1-3,6; and (4) fatty acid synthase (*Fasn*). We found significantly decreased expression for *Scd4* and *Elovl5* (**Figure 2B**) and an increased expression of *Fads2* and *Fasn* (**Figure 2B**). Other *Scd, Elvol,* and *Fads* isoforms were unchanged between WT (MyH6-Cas9^-^) and *Acly^KD^* mice (**Figure S3A**). *Scd4* is the rate-limiting enzyme in the biosynthesis of monounsaturated fatty acids (**Figure 2C**), while *Elovl5* is critical for the de novo synthesis of polyunsaturated fatty acids (PUFA). *Fads2* catalyzes the polyunsaturation of long-chain fatty acids, including 18:2(n-6), and its activity increases when *Scd* activity is reduced.^32, 33^ Our data indicate that loss of *Acly* shifts the degree of saturation from polyunsaturated fatty acids towards saturated shorter-chain fatty acids.

**Figure 2.**
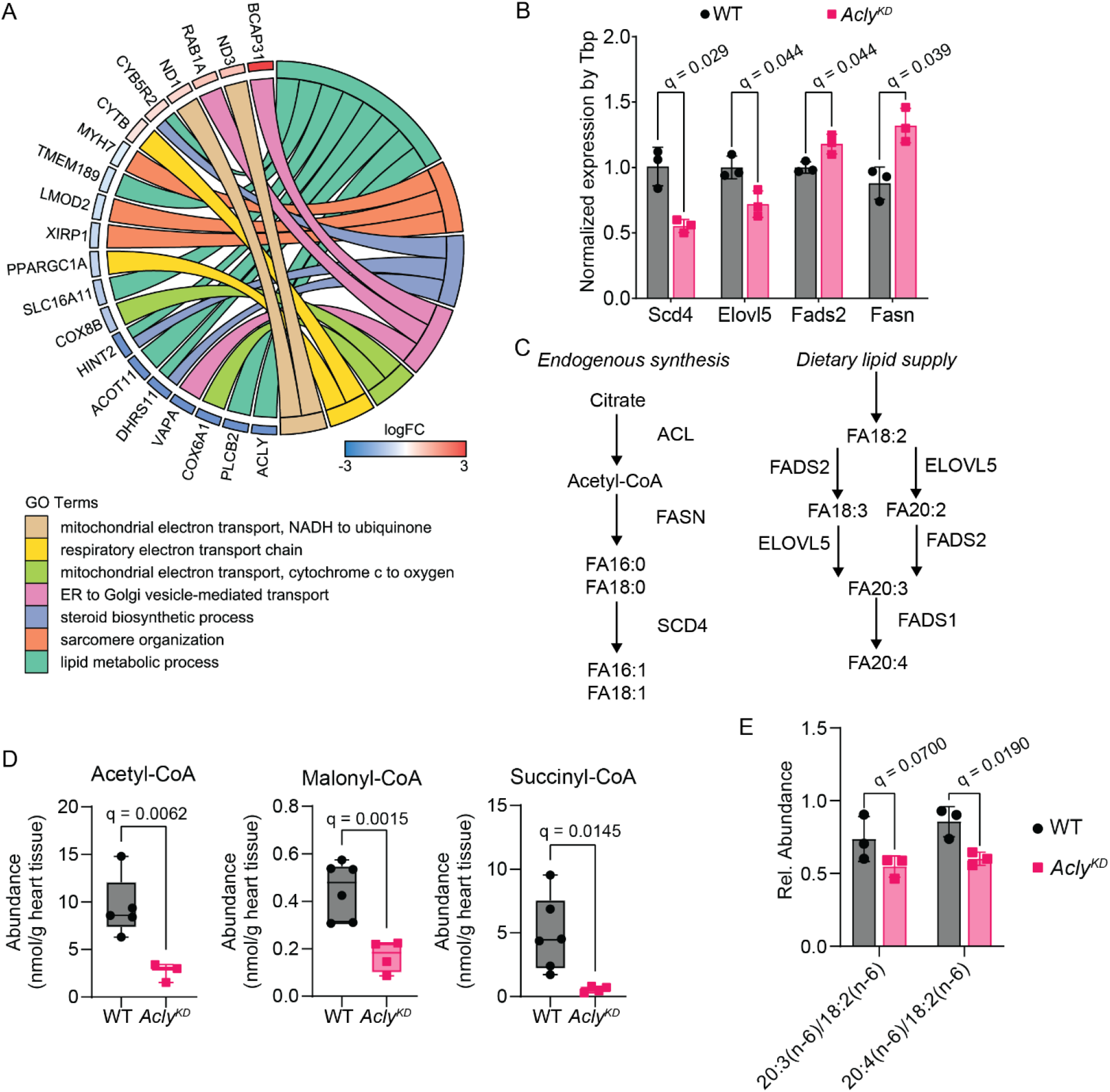
Functional analysis reveals increased lipid remodeling in *Acly^KD^* hearts. (A) Gene enrichment analysis of transcript changes from RNA-sequencing of adult mouse ventricular cardiomyocytes isolated from WT (MyH6-Cas9^-^) or *Acly^KD^* mouse hearts at 20 weeks. Genes with significant transcript changes and the top enriched gene ontology pathways are shown in the Chord diagram. Significant pathways are depicted on the right, and the fold change of core genes is shown on the left. Left-right connections indicate gene associations to pathways. (B) Gene expression of critical genes for fatty acid synthesis (*Fasn*), desaturation (*Fads2* and *Scd4*), and elongation (*Elovl5*). Expressions were quantified in heart tissue samples from WT (MyH6-Cas9^-^) or *Acly^KD^* mouse hearts at 20 weeks using RT-qPCR. Each biological sample was normalized by *Tbp* expression. n=4-6 male and female mice/group. 2-way ANOVA. Multiple comparison analysis by Sidak. (C) Schematic depicting interactions of fatty acid synthesis and modifications from endogenous and dietary sources. Abbreviations: *Elovl5*, fatty acid elongase 5; *Fads2*, fatty acid desaturase 2; *Fasn*, fatty acid synthase; *Scd4*, stearoyl-coenzyme A desaturase; *Tbp*, TATA-Box binding protein. (D) LC-MS/MS-based quantification of acetyl-, malonyl-, and succinyl-CoA in adult mouse ventricular cardiomyocytes isolated from WT (MyH6-Cas9^-^) or *Acly^KD^* mouse hearts at 20 weeks. n=4-6 male and female mice/group. 2-way ANOVA. Multiple comparison analysis by Sidak. (E) Relative abundance of polyunsaturated fatty acyl-esters using gas chromatography and mass spectrometry analysis in WT (MyH6-Cas9^-^) compared to *Acly^KD^* hearts at 20 weeks. The abundance of dihomo-gamma-linoleic acid (20:3) and arachidonic acid (20:4) was compared to linoleic acid (18:2). n=3 male and female mice/group. 2-way ANOVA. Multiple comparison analysis by Sidak. Numerical q-values are plotted for each comparison. Each data point represents a biological replicate, and error bars represent standard deviation.

To assess whether loss of ACL impairs precursors for endogenous lipid synthesis, we quantified acetyl-CoA, malonyl-CoA, and succinyl-CoA using targeted metabolomics by liquid chromatography and mass spectrometry (LC-MS/MS) profiling (see **Methods** for details). Total acetyl-CoA levels were decreased in *Acly^KD^* heart tissue compared to WT (MyH6-Cas9^-^) mice (**Figure 2D**). Surprisingly, we also detected decreased malonyl- and succinyl-CoA abundance in *Acly^KD^* heart tissue compared to WT (MyH6-Cas9^-^) mice (**Figure 2D**). The abundances of CoA species were unchanged in skeletal muscle (M. gastrocnemius), kidney, and lung tissue between control and *Acly^KD^* mice (**Figure S3B-D**). To further quantify the impact on lipid synthesis and modifications, we isolated membrane lipid fractions and quantified fatty acid compositions by tracing fatty acid methyl esters (FAMEs) using gas chromatography and mass spectrometry (GC-MS).^27, 34^ We found that cardiac-specific *Acly^KD^* increases the concentration of linoleic acid (LA; 18:2, n-6) compared to dihomo-gamma-linoleic acid (DHLA; 20:3, n-6) and arachidonic acid (20:4, n-6) causing an overall reduction in the DHLA to LA and AA to LA ratios (**Figure 2E**). These findings are consistent with the downregulation of Elovl5. Our data demonstrate that loss of *Acly* in the heart impacts both the *de novo* synthesis of fatty acids and overall fatty acid properties of membranes.

### *Acly^KD^* enhances glucose uptake *in vivo*

Because we observed significant up-regulation of mitochondrial complex I subunits at the mRNA levels, we hypothesized that *Acly^KD^* impacts energy substrate utilization in the heart. We used positron emission tomography/computer tomography (PET/CT) imaging combined with [^18^F]-Fluorodeoxyglucose ([^18^F]-FDG) to quantify myocardial glucose uptake *in vivo*. Tracer retention was normalized to plasma glucose level. We observed increased tracer retention in the heart (**Figure 3A**), corresponding to a 1.5-fold increased plasma glucose uptake in *Acly^KD^* mice compared to WT (MyH6-Cas9^-^) littermates (**Figure 3B** and **Figure S3E**). Changes in glucose utilization were limited to cardiac tissue and not observed in the kidneys (**Figure 3B** and **Figure S3E**). We did not observe a significant difference in plasma glucose levels between experimental groups (**Figure 3C**), suggesting that metabolic alterations in response to *Acly* deficiency increase [^18^F]-FDG uptake. To corroborate these findings, we quantified metabolite levels and enzyme activities from heart tissue samples. Compared to WT littermates, *Acly^KD^* mice exhibited lower ATP levels (**Figure 3C**) in the heart. *Acly^KD^* mice displayed reduced ACL activity (**Figure 3D**) but increased PFK activity (**Figure 3E**) and increased glucose 6-phosphate levels (**Figure 3F**). These data indicate that although ACL regulates acetyl-CoA synthesis, this enzyme is required for cardiac energy homeostasis, and loss increases glucose uptake and glycolytic flux in the heart.

**Figure 3.**
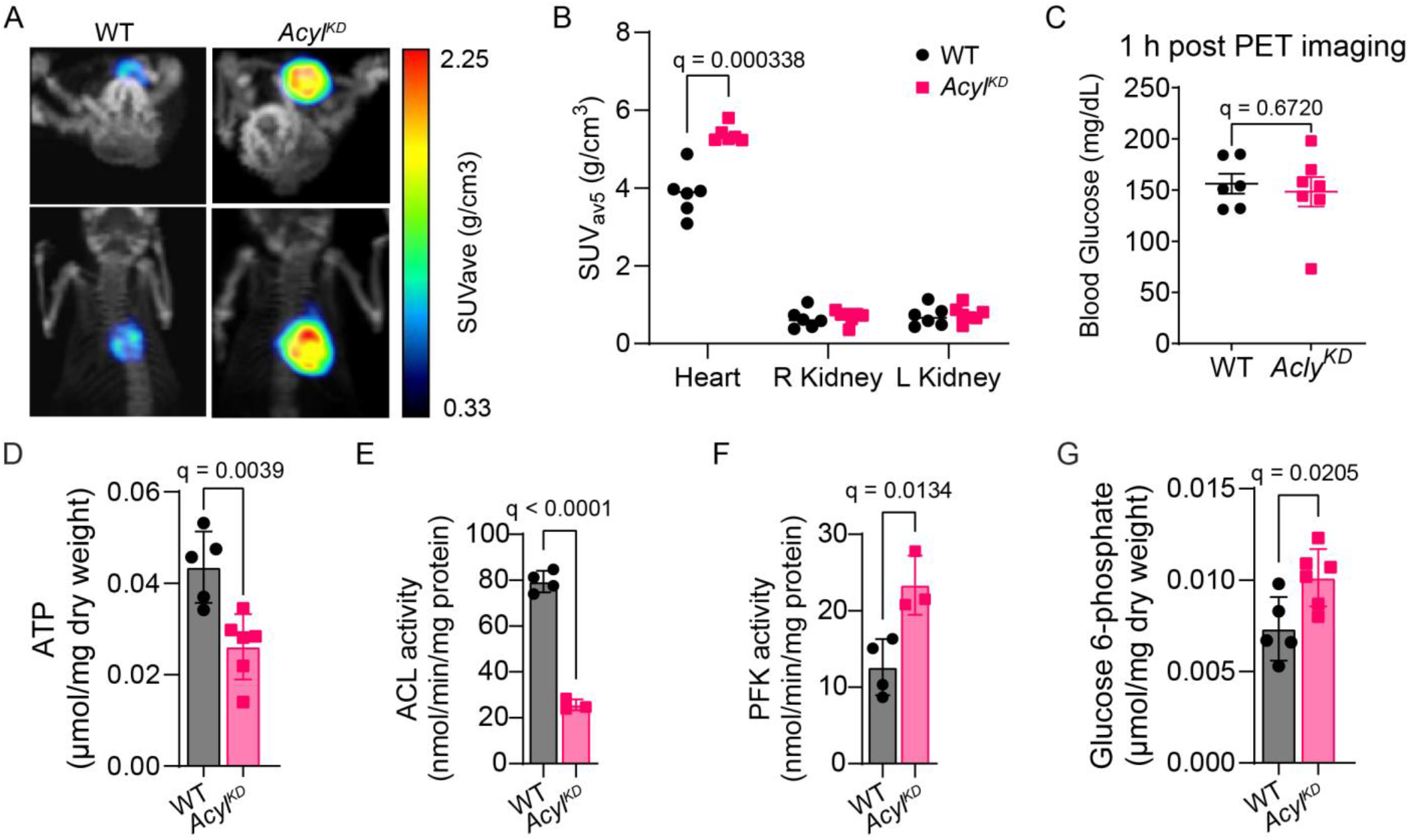
Loss of *Acly* promotes increased glucose oxidation *in vivo*. (A) Representative images of [^18^F]-fluorodeoxyglucose (FDG)-positron emission tomography analysis *in vivo* of WT (MyH6-Cas9^-^) or *Acly^KD^* (MyH6-Cas9^+^) mice at 20 weeks. (B-C) Tracer retentions (B) relative to weight-normalized injected dose (SUV) in heart and kidneys *of in vivo*-injected WT (MyH6-Cas9^-^) or *Acly^KD^* (MyH6-Cas9^+^) mice at 20 weeks. The highest five values were used in the average. SUVs were normalized to plasma glucose levels (C). n=7 male and female mice/group. 2-way ANOVA. Multiple comparison analysis by Sidak. FDR<1%. (D-E) Enzymatic activities for ATP-dependent citrate lyase (ACL) (D) and Phosphofructokinase (PFK) (E) were quantified in heart tissue samples from WT (MyH6-Cas9^-^) or *Acly^KD^* (MyH6-Cas9^+^) mice at 20 weeks using enzymatic assays. n=4 male and female mice/group. 2-way ANOVA. Multiple comparison analysis by Sidak. FDR<1%. (F-G) Glucose-6 phosphate (F) and ATP (G) levels were quantified in heart tissue samples from WT (MyH6-Cas9^-^) or *Acly^KD^* (MyH6-Cas9^+^) mice at 20 weeks using spectrophotometric assays. n=5-7 male and female mice/group. 2-way ANOVA. Multiple comparison analysis by Sidak. FDR<1%. Numerical q-values are plotted for each comparison. Each data point represents a biological replicate, and error bars represent standard deviation.

### *Acly* deficiency impairs cardiac oxidative metabolism

To investigate how the loss of *Acly* impacts cardiac oxidative metabolism, we conducted heart perfusions with stable isotope tracer labeling followed by targeted GC-MS-based metabolomics and metabolic flux analysis. Hearts from *Acly^KD^* and WT (MyH6-Cas9^-^) mice were perfused *ex vivo* for 30 min using the working heart preparation (**Figure 4A**). We observed a decreased contractile function (**Figure 4B**) and myocardial oxygen consumption (**Figure 4C**) in *Acly^KD^* hearts. The metabolic fate of glucose and palmitate was determined using ^13^C_6_-glucose or ^13^C_16_-palmitate in the presence of physiological concentrations of unlabeled lactate, pyruvate, and glutamine (**Figure 4D**; see Methods for details). We observed decreased lactate m+3 labeling and m+2 labeling in Krebs cycle intermediates (e.g., citrate, malate, and fumarate) (**Figure 4E**). These findings are consistent with a decreased oxidative decarboxylation of glucose and increased release of lactate into the coronary effluent during the perfusion. The citrate m+2 to malate m+2 ratio was significantly decreased in *Acly^KD^* compared to WT (MyH6-Cas9^-^) littermates. The reduced transition of citrate m+2 toward malate m+2 indicates that loss of *Acly* in cardiomyocytes reduces oxidative metabolism in the Krebs cycle. Using LC-MS/MS, we quantified fatty acid β-oxidation of palmitate by measuring ^13^C_16_-palmitoylcarnitine in tissue samples from perfused hearts. Loss of *Acly* reduced ^13^C_16_-palmitoylcarnitine levels in heart tissue by 3-fold (**Figure 4F**). These data indicate enhanced glycolysis compared to fatty acid oxidation to provide ATP when ACL activity is low, consistent with an impaired mitochondrial function and oxidative phosphorylation.

**Figure 4.**
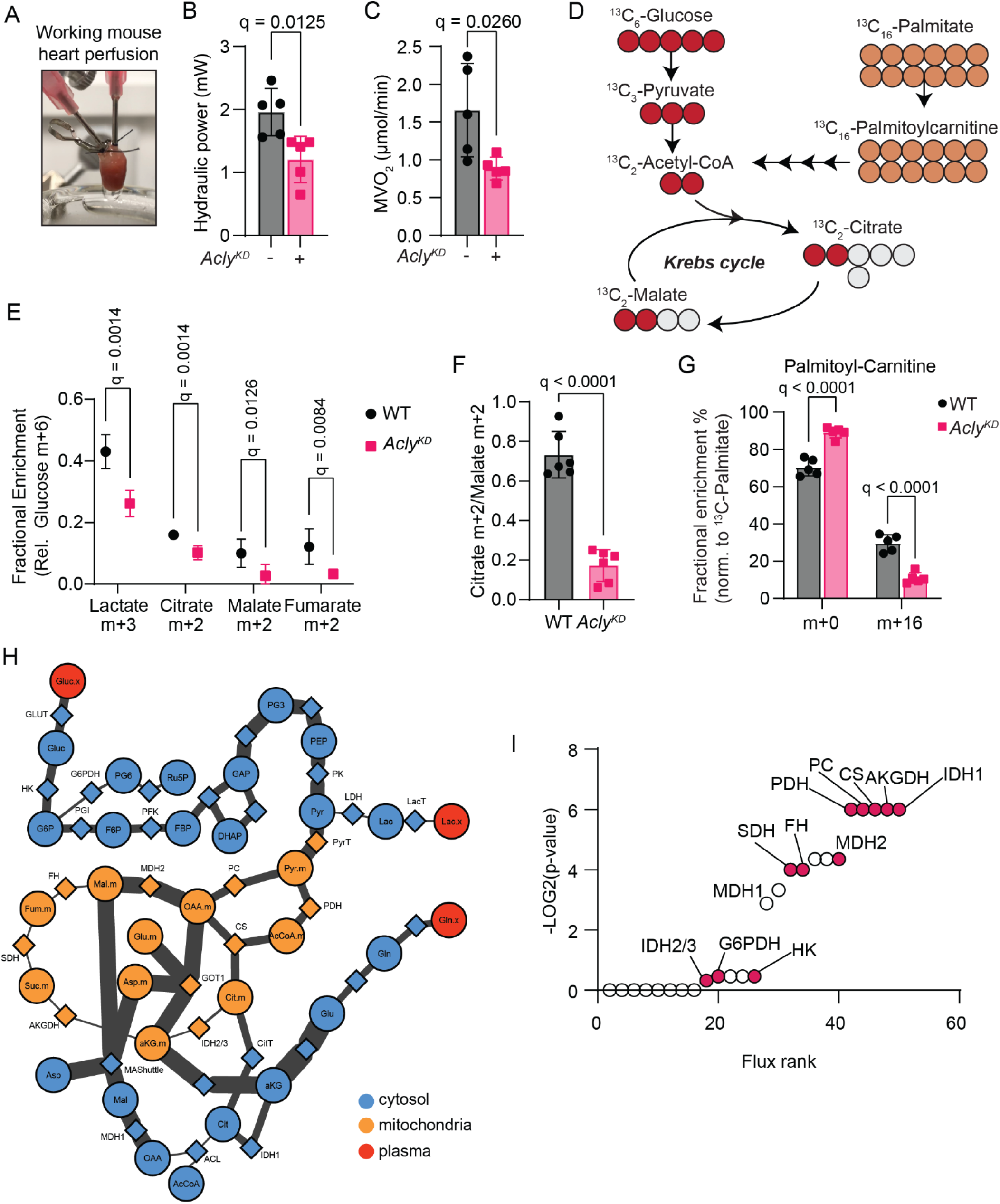
Loss of *Acly* shifts glucose and palmitate contribution to oxidative metabolism in the heart. (A) Experimental set-up of working mouse heart perfusions. (B-C) Hydraulic power (B) and myocardial oxygen consumption (MVO_2_) (C) in perfused working mouse hearts from WT (MyH6-Cas9^-^) or *Acly^KD^* male and female mice at 20 weeks. (D) Schematic of the theoretical isotopomer distribution in central carbon metabolism using ^13^C_6_-glucose and ^13^C_16_-palmitate tracers. Metabolites are indicated in black circles (representing carbons). Filled red circles and organ circles refer to heavy carbons (^13^C). (E) Fractional enrichment of the lactate m+3, citrate m+2, malate m+2, and fumarate m+2 from steady-state ^13^C_6_-glucose perfusions of WT (MyH6-Cas9^-^) or *Acly^KD^* mouse hearts at 20 weeks. n = 6 male and female mice/group. Statistical significance was calculated by multiple unpaired t-tests followed by multiple comparisons analysis using a false discovery rate < 1% by the two-step method of Benjamini, Krieger, and Yekutieli. Adjusted q-value is blotted for each comparison. (F) Malate m+2 to Citrate m+2 ratio indicates a shift in ^13^C_6_-glucose-derived carbon distribution in WT compared to *Acly^KD^* mouse hearts. n = 6 male and female mice/group. Statistical significance was calculated by multiple unpaired t-tests followed by multiple comparisons analysis using a false discovery rate < 1% by the two-step method of Benjamini, Krieger, and Yekutieli. Adjusted q-value is blotted for each comparison. (G) Fractional enrichment of palmitoyl-carnitine in ^13^C_16_-palmitate working mouse heart perfusions. n = 5 male and female mice/group. Statistical significance was calculated by multiple unpaired t-tests followed by multiple comparisons analysis using a false discovery rate < 1% by the two-step method of Benjamini, Krieger, and Yekutieli. Adjusted q-value is blotted for each comparison. (H) Comparative metabolic flux analysis using ^13^C_6_-glucose isotopomer distributions in WT (MyH6-Cas9^-^) or *Acly^KD^* mouse hearts. Isotopologues from each experimental group were fitted to the depicted fully compartmentalized network. Metabolites and enzymes are depicted as circles and diamonds, respectively. Carbon transitions were assigned to cytosolic, mitochondrial, or extracellular metabolic fluxes. The thickness of each line connecting metabolites and proteins indicates the estimated flux rate in *Acly^KD^* compared to WT. (I) Calculated flux rates were statistically compared between WT (MyH6-Cas9^-^) and *Acly^KD^* groups. Enzymes were ranked according to their associated flux rate change and log2(p-value). Abbreviations: AKGDH, alpha-ketoglutarate dehydrogenase; CS, citrate synthase; FH, fumarate hydratase; G6PDH, glucose-6 phosphate dehydrogenase; HK, hexokinase; IDH, isocitrate dehydrogenase; MDH, malate dehydrogenase; PC, pyruvate carboxylase; PDH, pyruvate dehydrogenase; SDH, succinate dehydrogenase.

Next, we integrated the isotopomer distribution data into metabolic flux analysis (MFA) and conducted computational simulations to determine theoretical flux distributions in our model^5, 30^. We generated a simplified network of carbon transitions, including glycolysis and Krebs cycle, to reflect our experimental dataset (see **Table 2**). *Acly^KD^* simulations displayed a high glycolytic flux and increased release of lactate (**Figure 4H; Supplemental Table 2** and **Supplemental Table 3**). The model predicted an increased contribution of malate and aspartate shuttling between mitochondria and cytosol to compensate for the absence of ACL. Furthermore, increased flux through the cytosolic and NADPH-dependent isocitrate dehydrogenase (IDH) 1 abrogated citrate accumulation in the cytosol, suggesting that *Acly^KD^* increases the NADPH redox state (**Figure 4H** and **Table 3**). We ranked flux rates to identify which enzymes drive metabolic adaptation in response to the loss of Acly. Surprisingly, IDH1 was the top altered enzyme in *Acly^KD^* compared to WT (MyH6-Cas9^-^) controls (**Figure 4I**). These data demonstrate that ACL deficiency shifts fatty acid oxidation towards glucose and increases cytosolic IDH1 flux to regulate citrate levels in the cytosol.

### Metabolic flux changes are regulated by YAP in *Acly^KD^* hearts

MFA raised the question of how glycolysis and citrate metabolism are upregulated in the heart under conditions of *Acly* loss and reduced mitochondrial oxidative metabolism. To address this question, we quantified the heart tissue expression of IDH1, 2, and 3 using western blotting. Tissue samples from *Acly^KD^* mice displayed an increased IDH1 expression compared to WT (MyH6-Cas9^-^) controls, while IDH3 expression was decreased (**Figure 5A** and **B**), suggesting a redirection of cytosolic citrate into α-KG. Pathway analysis of RNA-seq data revealed a downregulation and inhibition of TEAD1, which interacts with the YAP (Yes-associated protein) and TAZ (transcriptional coactivator with PDZ-binding motif) complex. YAP/TAZ expression was significantly downregulated by western blotting in *Acly^KD^* hearts compared to WT littermates (**Figure 5C** and **D**). Next, we measured the phosphorylation of YAP/TAZ in serine residues 127 and 397. Phosphorylation of YAP/TAZ was significantly increased, especially at S397 (**Figure 5E** and **F**). Increased phosphorylation of YAP/TAZ promotes their cytoplasmic localization through binding to 14-3-3 proteins and degradation, resulting in decreased transcriptional activation at the TEA domain.

**Figure 5.**
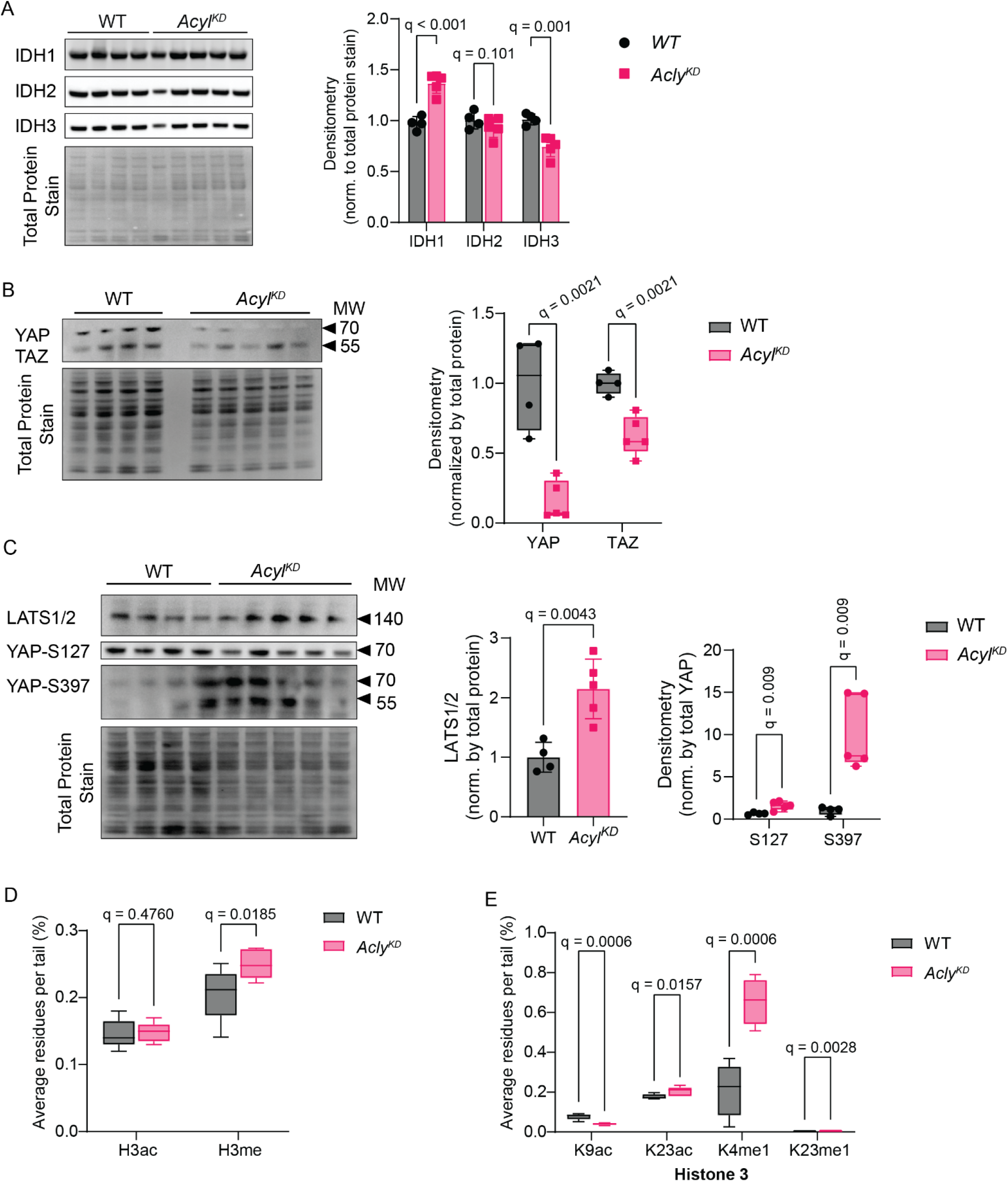
Loss of *Acly* induces overexpression of IDH1 and dephosphorylation of YAP. (A-B) Protein expression of isocitrate dehydrogenase (IDH) 1, 2, and 3 isoforms in heart tissue samples from WT (MyH6-Cas9^-^) and *Acly^KD^* hearts at 20 weeks. Representative western blot images with total protein stains (A) and densitometry (B). n=4-5 male and female mice/group. 2-way ANOVA. Multiple comparison analysis by Sidak. FDR<5%; adjusted q-values are depicted on each bar plot. (C-D) Representative western blot images (C) and densitometry (D) of total YAP expression in WT (MyH6-Cas9^-^) and *Acly^KD^* hearts at 20 weeks. n=4-5 male and female mice/group. 2-way ANOVA. Multiple comparison analysis by Sidak. FDR<5%; adjusted q-values are depicted on each bar plot. (E-F) Representative western blot images (C) and densitometry (D) of YAP phosphorylation at serine (S) residues 127 and 397 in WT (MyH6-Cas9^-^) and *Acly^KD^* hearts. n=4-5 male and female mice/group. 2-way ANOVA. Multiple comparison analysis by Sidak. FDR<5%; adjusted q-values are depicted on each bar plot.

We considered the possibility that reduced transcriptional activation by YAP/TAZ and increased availability of α-KG by IDH1 induce heterochromatin structures marked by H3 acetylation and methylation changes. We isolated histones and quantified histone 3 acetylation and methylation using MS-based proteomics. YAP/TAZ depletion and *Acly^KD^* were associated with increased histone 3 pan-methylation (**Figure 5D**) and a marked reduction in H3K9ac (**Figure 5E**), while acetylation at K14, K18, K27, K56, and K79 was unaffected (**Figure S5A**). Intriguingly, we identified an increased H3K23 acetylation in *Acly^KD^* hearts compared to control hearts (**Figure 5E**). We identified an increased mono-methylation at H3K4 and H3K23 (**Figure 5E**), while di- and trimethylation at other histone residues was unaffected (**Figure S5B** and **S5C**). Our findings indicate that YAP/TAZ depletion activates gene expression via histone 3 acetylation, while ACSS2 maintains acyl-CoA synthesis without ACL.

### *In silico* blockage of IDH1 flux stimulates oxidative metabolism

Next, we explored how cardiomyocytes can overcome the loss of ACL using computational modeling with CardioNet.^5, 7, 21, 29, 30^ We annotated our gene expression, enzyme activities, and metabolite changes to CardioNet (see **Methods** for details). We determined flux distributions using flux balance analysis (FBA), which applies a steady-state assumption while optimizing an objective function. The linear optimization problem was to maximize ATP hydrolysis and biomass synthesis while fulfilling experimental constraints defined by the working mouse perfusions of WT and *Acly^KD^* hearts. Annotation enrichment of significantly altered flux rates (**Figure 6A**) demonstrated distinct metabolic remodeling between WT (MyH6-Cas9^-^) controls and *Acly^KD^* hearts consistent with our multi-omics analysis. Consistent with our tracer-based studies, CardioNet simulations revealed that loss of ACL triggers an upregulation of glucose metabolism. In addition, the modeling revealed increased contribution of amino acid metabolism, including glutamate metabolism, while oxidative phosphorylation and nucleotide metabolism are impaired. To identify alternative metabolic pathways compensating for the loss of ACL, we tasked the model to achieve ATP hydrolysis at the WT level while using constraints from *Acly^KD^* simulations. PCA analysis (**Figure 6B**) revealed that these *Acly^Rescue^* simulations attenuated the metabolic disruption caused by *Acly^KD^*. We identified key-regulatory enzymes within oxidative and reductive (**Figure S6A**) metabolic pathways that can normalize flux rates to the WT level. Further inspection of the flux distribution between *Acly^Rescue^* and *Acly^KD^* simulations revealed a subset of 22 candidate enzymes (**Figure 6C**). We conducted *in silico* knockout simulations for each candidate and tested their efficacy in reversing the metabolic adaptation caused by loss of ACL. We simulate an enzymatic function for each candidate enzyme from 0% to 100%. Our simulations identified seven candidate enzymes that increased oxidative phosphorylation (OXPHOS) (**Figure 6D** and **Figure S6B**). Only GAPDH and IDH1 knockout simulations reduced IDH1 flux and improved OXPHOS and nucleotide metabolism (G6PD), indicative of augmented metabolic function. However, a direct comparison of simulations revealed that blocking IDH1 progressively increases OXPHOS (**Figure 6E**) and the mitochondrial NADH redox potential (**Figure S6C**). Our simulations suggest that IDH1 is a potential metabolic vulnerability in *Acly^KD^* hearts.

**Figure 6.**
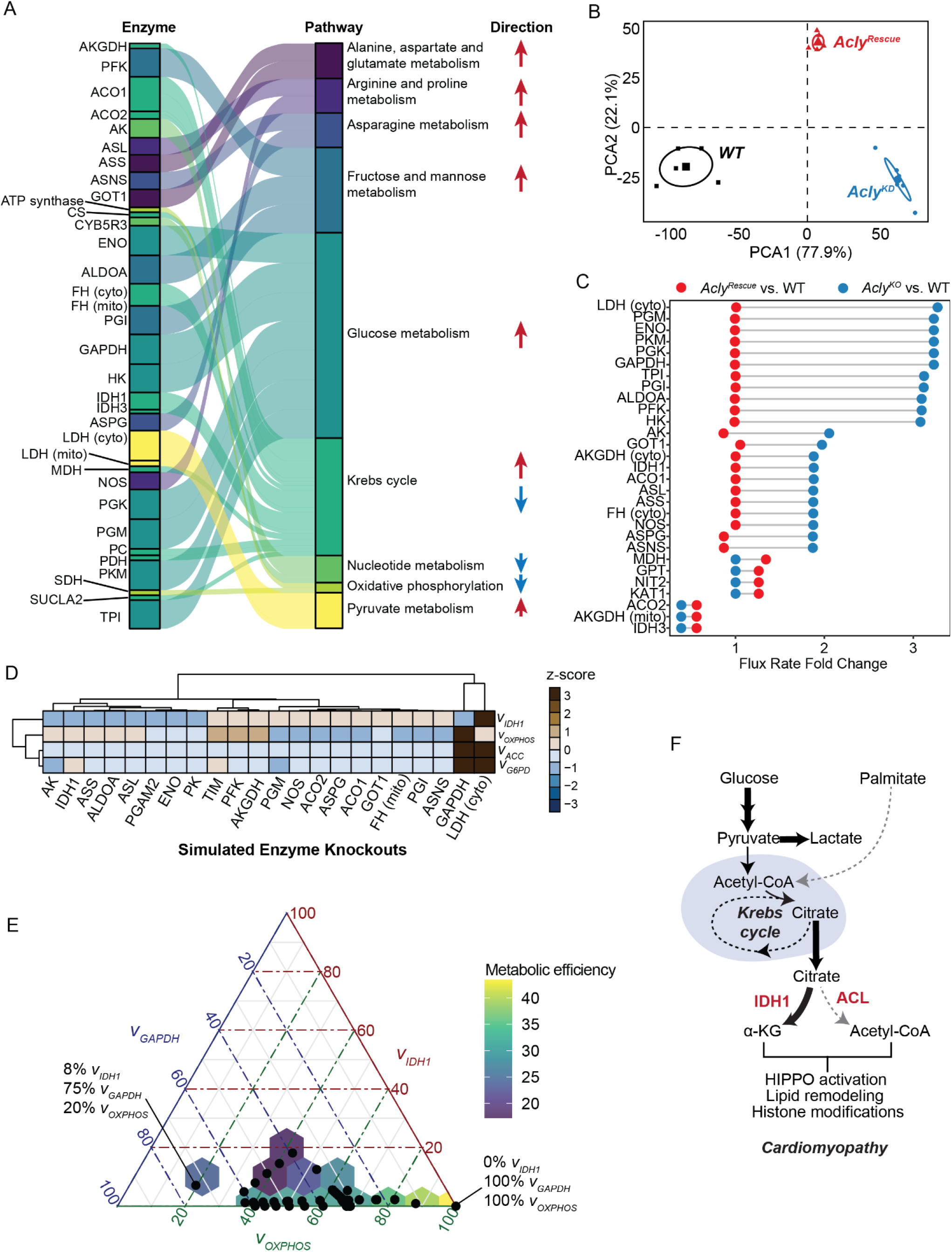
CardioNet simulation reveals metabolic vulnerabilities in *Acly^KD^*. (A) Regression analysis of computational flux simulations in WT (MyH6-Cas9^-^) and *Acly^KD^* hearts. Experimental data from working mouse heart perfusions was integrated into CardioNet simulations, and metabolic flux rates were calculated using an objective function. Calculated flux rates were compared using a linear regression model and annotated to CardioNet pathways. (B) Principal component analysis (PCA) of calculated flux distributions from CardioNet simulations of experimental WT (MyH6-Cas9^-^), *Acly^KD^*, and theoretical *Acly^Rescue^* hearts. PCA components 1 and 2 describe 100% of the variability in the simulations. (C) Differences in flux rate fold changes of significantly altered metabolic reactions between *Acly^Rescue^* and *Acly^KD^* simulations compared to WT simulations. Indicated enzymes are potential metabolic vulnerabilities in response to loss of ACL. (D) Comparison of critical metabolic fluxes in simulated enzyme knockouts using CardioNet. Theoretical flux distributions from each knockout simulation were compared to WT simulations according to the calculated *vIDH1*, *vOXPHOS*, *vACC*, and *vG6PD*. (E) Ternary plot depicting the distribution and variability of GAPDH and IDH1 knockout simulations as a function of *vIDH1*, *vGAPDH,* and *vOXPHOS.* Each side of the ternary plot represents one of the three components: *vIDH1*, *vGAPDH,* or *vOXPHOS*. Each point in the ternary plot corresponds to a value that intersects perpendicular to each leg of the triangle. Examples are depicted for two simulations. (F) Schematic of metabolic adaptation in response to loss of Acly and metabolic drivers for cardiac remodeling. Reduction of ACL causes increased cytosolic IDH1 flux into α-KG and reduced de novo synthesis of acetyl-CoA, which act as a metabolic signal promoting HIPPO activation and lipid remodeling histone modifications, ultimately driving the development of cardiomyopathy. Abbreviations: ACO, aconitase; ACC, acetyl-coA carboxylase; ALDOA, aldolase A; AK, adenylate kinase; AKGDH, alpha-ketoglutarate dehydrogenase; ASL, Argininosuccinate lyase; ASS, Argininosuccinate synthase; ASNS, Asparagine synthetase; ASPG, L-asparaginase; CS, citrate synthase; CYB5R3, NADH-cytochrome b5 reductase 3; ENO, enolase; FH, fumarate hydratase; GOT1, glutamic-oxaloacetic transaminase 1; GAPDH, Glyceraldehyde 3-phosphate dehydrogenase; G6PDH, glucose-6 phosphate dehydrogenase; GPT, Alanine aminotransferase; HK, hexokinase; IDH, isocitrate dehydrogenase; MDH, malate dehydrogenase; NOS, nitric oxide synthase; OXPHOS, oxidative phosphodrylation; PC, pyruvate carboxylase; PDH, pyruvate dehydrogenase; PGI, phosphoglycerate isomerase; PGK, Phosphoglycerate kinase; PGM, Phosphoglucomutase; PGAM, Phosphoglycerate mutase; PKM, pyruvate kinase; PFK, phosphofructokinase; SDH, succinate dehydrogenase; SUCLA; Succinate--CoA ligase, TPI, Triosephosphate isomerase.

## DISCUSSION

Our multi-omics analysis revealed an unexpected metabolic finding: ACL activity is critical for cardiac contractile function and modulates lipid synthesis in the heart. This result contrasts the well-accepted perception of cardiac metabolism to rely on exogenous lipid sources. Our study expands the current understanding of oxidative and reductive metabolism with enhanced glycolysis without mitochondrial oxidative metabolism. Heart failure is characterized by increased glucose uptake and decreased fatty acid oxidation, interpreted as deficient mitochondrial oxidative phosphorylation. However, our findings indicate that oxidative decarboxylation of nutrients supports reductive processes and ensures the production of reducing equivalents in the heart during metabolic stress. ACL is required in the heart to sustain ATP and citrate homeostasis. Our data shows that cardiac metabolism requires de novo acetyl-CoA synthesis and can synthesize fatty acid under normal physiologic conditions.

Increased glucose uptake and oxidation serve multiple purposes. Our data support the established concept that the heart shifts from one energy-providing substrate towards another during stress. These findings are consistent with maladaptive metabolic remodeling and reversal to the fetal gene program described previously in the failing and hypertrophic heart.^1, 5, 35^ Our results indicate that YAP/TAZ signaling supports metabolic adaptation during impaired ACL function. Several YAP/TAZ-regulated transcripts are down-regulated in our animal model. Reduced transcriptional recruitment of YAP/TAZ to the chromatin is associated with increased histone 3 acetylation.^36^ We observed increased H3K23 acetylation, supporting gene activation and cardiac adaptation.

Metabolic adaptation is driven by transcriptional and allosteric regulation of enzymes. A surprising role for ACL in the heart is the increased control of glucose utilization via regulation of metabolic flux. Our data demonstrate an increased decarboxylation of citrate during loss of ACL, suggesting that a substantial portion of citrate is exported during wildtype conditions. Citrate regulates glycolysis through allosteric inhibition of PFK when glucose uptake exceeds oxidative capacity. Our findings support this concept and demonstrate an increased citrate abundance during loss of myocardial ACL. However, our data support increased oxidation of glucose-derived citrate, which is partially regulated by NADPH-dependent isocitrate dehydrogenases and non-canonical functions of the Krebs cycle.

The loss of ACL in the heart induces fatty acyl-chain modifications and changes in structural lipids, an unexpected change that impacts cellular membrane integrity and homeostasis. Although acetyl-CoA is a critical intermediate of the Krebs cycle during oxidative metabolism, it has been appreciated as a precursor for de novo fatty acid and cholesterol synthesis in the heart. ^18, 37^ The precise contribution of endogenous acetyl-CoA synthesis in the heart under normal physiological conditions is still not fully understood. *In vivo* studies are further complicated by complicated tracer patterns due to liver and kidney metabolism interferences. Our approach to combining ex vivo tracer studies with computational modeling aims to overcome these shortcomings and provide a holistic understanding of cardiac metabolic adaptation. CardioNet simulations predict an increased contribution of pentose phosphate pathway flux to produce NAPDH from glucose. Several key regulatory enzymes within the lipid metabolism require NAPDH as a precursor, which will have activity-dependent changes. These compensatory flux changes likely ensure cardiac adaptation during the loss of ACL function and support the increased import of glucose and amino acids for alternative fates rather than solely ATP provision, including protein synthesis. Our findings emphasize the role of ACL and IDH1 in regulating cardiac energy substrate metabolism. Therefore, therapeutic strategies targeting ACL must consider metabolic adaptation in the heart and the potential for cardiomyopathy in patients.

## ACKNOWLEDGEMENTS

We thank the Histopathology Laboratory, Department of Pathology and Laboratory Medicine, at McGovern Medical School at The University of Texas Health Science Center at Houston for generating histology samples. We thank the IDDRC Neuroconnectivity Core at Baylor College of Medicine (Houston, TX) for virus packaging. Figures were created with BioRender.com.

## AUTHOR CONTRIBUTIONS

Conceptualization, A.K.; Methodology, A.K., J.F.M., S.L., S.T.G., D.P.W.; Metabolomics and FAMEs, A.K., L.T., K.K., E.C.L., P.L.L., M.G.V.D.H.; PET-CT, S.T.G., D.P.W.; Proteomics, F.N.V., J.M.G., B.A.G.; Computational Modeling, A.K., K.K.; RNA-sequencing, B.M.H., A.Q.D.; Data analysis, S.L., A.K., E.C.L., S.T.G.; Investigation, S.L., B.D.G., Y.G., K.K., I.W., J.P., A.D., R.K., B.D.G., R.L.S.; Writing – Original Draft, S.L. and A.K.; Writing – Review & Editing, S.T.G., L.T., Y.G., K.K., I.W., J.P., A.D., R.K., B.D.G., R.L.S., A.Q.D., E.C.L., F.N.V., J.M.G., B.M.H, B.A.G., M.G.V.H., P.L.L., H.T., D.P.W., J.F.M.; Resources, J.F.M., H.T., D.P.W., P.L.L., B.A.G, M.G.V.H.

