## Supplemental Materials for "ATP-dependent citrate lyase Drives Left Ventricular Dysfunction by Metabolic Remodeling of the Heart"

**TITLE**

**AFFILIATIONS**

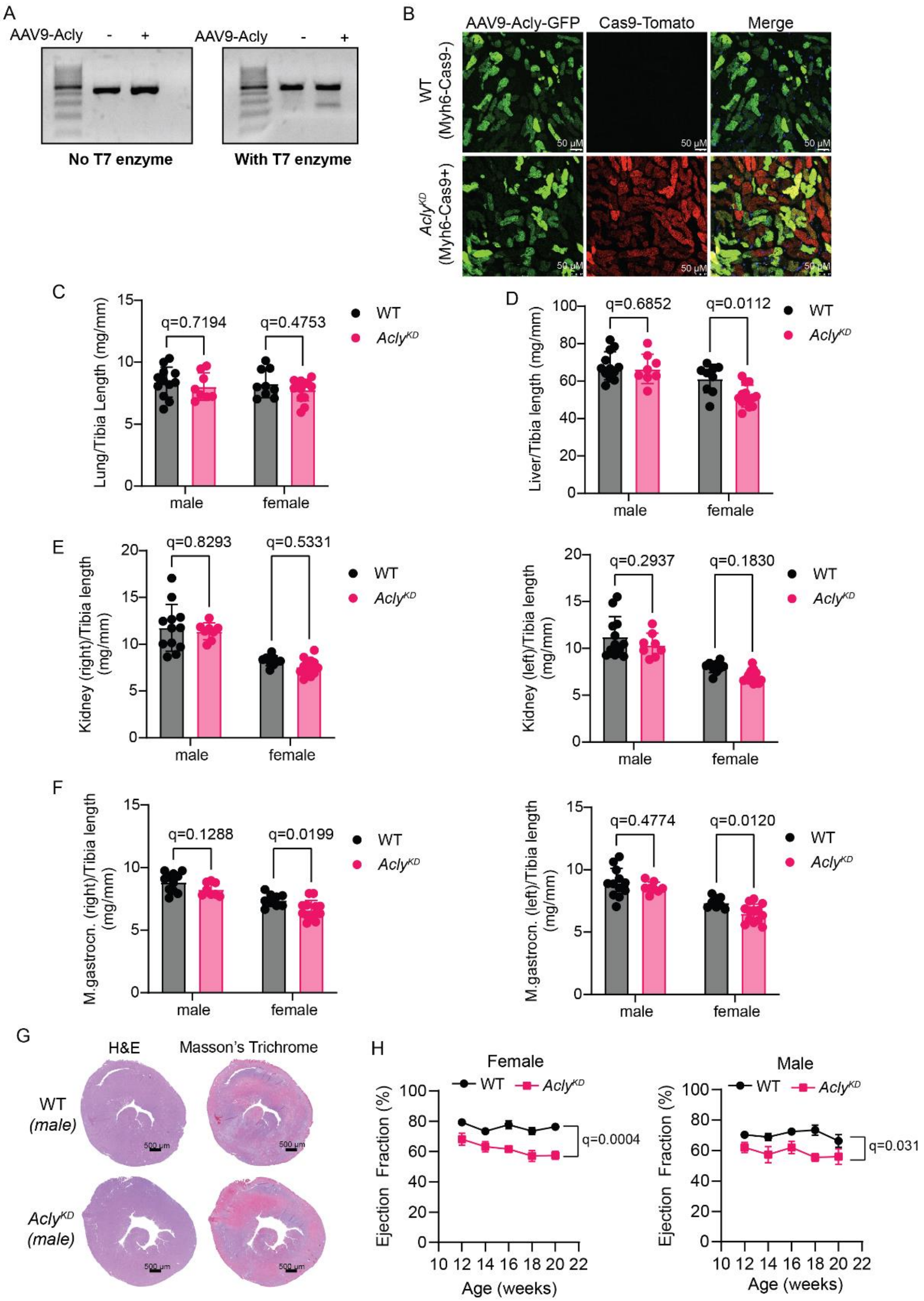


**Supplemental Figure 1. *Acly*-knockdown impairs cardiac function.**
(A) AAV9-sgRNAs targeting *Acly* were evaluated by double-strand break induction efficiency using T7 endonuclease assays (T7 enzyme) in adult mouse ventricular cardiomyocytes (AMVMs) isolated from MyH6-Cas9$^+$ mice. Representative images of T7 endonuclease assays demonstrate the double-strand break in response to AAV9-gRNA transduction.

(B) Representative images of $Cas9^{+}$ expression and Acly-knockdown efficiency in AMVMs isolated from MyH6-$Cas9^{-}$ (WT) and MyH6-$Cas9^{+}$ ($Acly^{KD}$) mice. scale bar equals 50 µm.
(C-F) Organ weight (mg) normalized to tibia length (mm) from WT (MyH6-$Cas9^{-}$) and $Acly^{KD}$ (MyH6-$Cas9^{+}$) mice. Lung (E), liver (F), kidneys (G), and M. gastrocnemius (H) were compared from male and female WT and $Acly^{KD}$ mice. n=10-14 male and female mice/group. 2-way ANOVA. Multiple comparison analysis by Sidak. Numerical q-values are plotted for each comparison. Each data point represents a biological replicate, and error bars represent standard deviation.
(G) Representative images of H&E and Masson's trichrome staining from male and female WT (MyH6-$Cas9^{-}$) and $Acly^{KD}$ mouse hearts.
(H) Cardiac function was evaluated by echocardiography at 12, 14, 16,18, and 20 weeks in male and female WT (MyH6-$Cas9^{-}$) and $Acly^{KD}$ mice. n = 7 male and female mice/group. Statistical significance was calculated by multiple unpaired t-tests followed by multiple comparisons analysis using a false discovery rate < 5% by the two-step method of Benjamini, Krieger, and Yekutieli.

| | **WT, N= 7** | ***Acly*$^{KD}$, N=7** | **p-value** | **q-value** |
|---|---|---|---|---|
| Heart Rate (BPM) | 522.9 (± 40.95) | 485.6 (± 42.99) | 0.1224 | 0.0466 |
| Diameter;systolic (mm) | 2.127 (± 0.27) | 2.914 (± 0.48) | **0.0028** | **0.0034** |
| Diameter;diastolic (mm) | 3.501 (± 0.18) | 4.079 (± 0.4) | **0.0044** | **0.0034** |
| Volume;systolic (μL) | 15.35 (± 4.97) | 34.02 (± 14.47) | **0.0073** | **0.0037** |
| Volume;diastolic (μL) | 51.13 (± 6.27) | 74.26 (± 17.54) | **0.0065** | **0.0037** |
| Stroke Volume (μL) | 35.78 (± 3.82) | 40.24 (± 5.72) | 0.1118 | 0.0466 |
| Ejection Fraction (%) | 70.45 (± 6.97) | 55.64 (± 8.73) | **0.0043** | **0.0034** |
| Fractional Shortening (%) | 39.4 (± 5.68) | 28.92 (± 5.57) | **0.0045** | **0.0034** |
| Cardiac Output (mL/min) | 18.66 (± 1.92) | 19.56 (± 3.4) | 0.5496 | 0.1859 |

**Supplemental Table 1. Evaluation of cardiac function using echocardiography at 20 weeks in WT (MyH6-Cas9$^{-}$) and *Acly*$^{KD}$ (MyH6-Cas9$^{+}$) mice.** Echocardiographic imaging for control and *Acly*$^{KD}$ mice was analyzed between 14 to 20 weeks of age. Unpaired t-test; significance threshold: p-value<0.05 and q-value FDR<1% (two-stage-up method of Benjamini, Krieger, and Yekutieli).

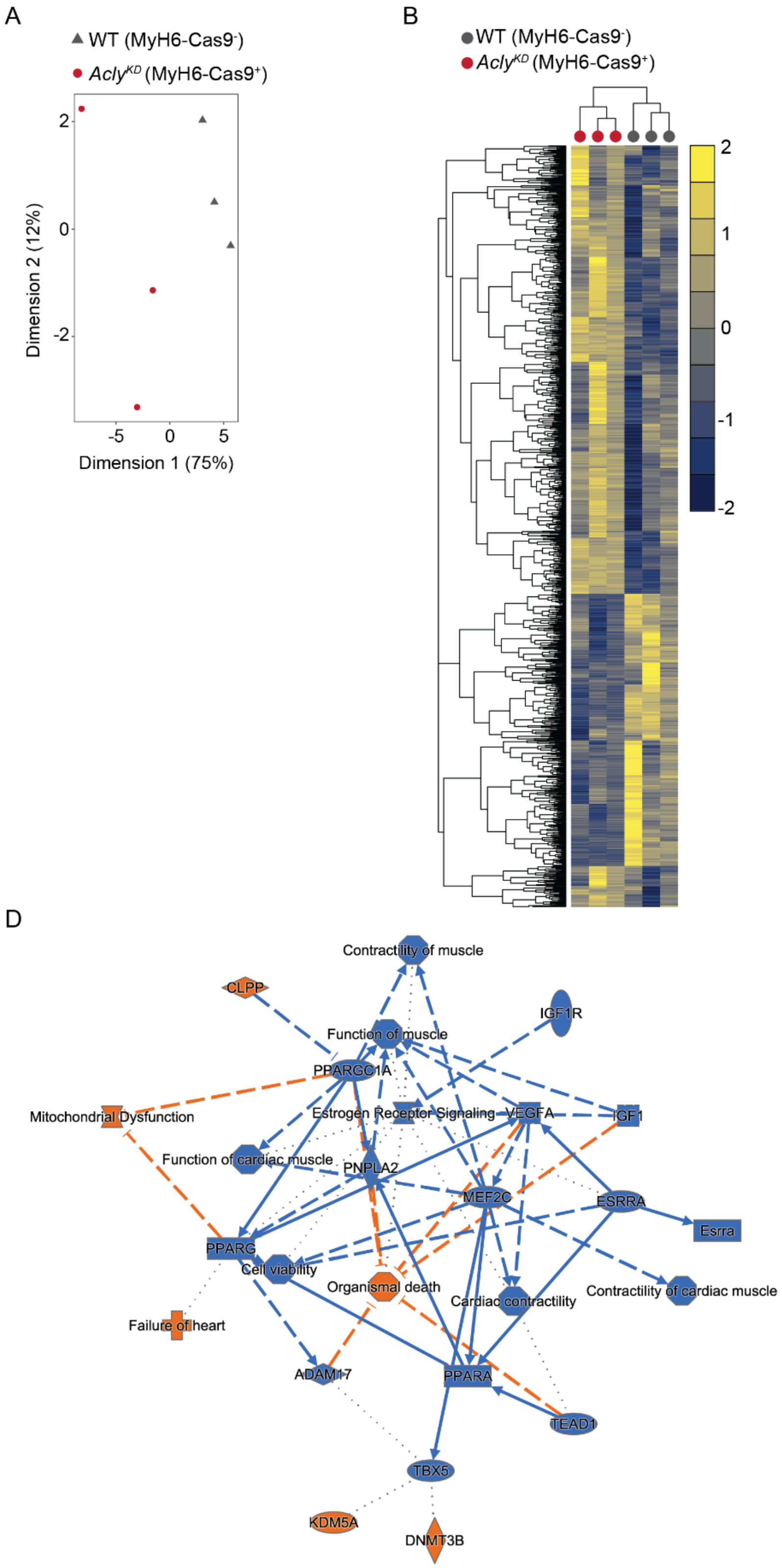
A
WT (MyH6-Cas9-)
Acly^KD (MyH6-Cas9+)
Dimension 2 (12%)
Dimension 1 (75%)
2
0
-2
-5
0
5
B
WT (MyH6-Cas9-)
Acly^KD (MyH6-Cas9+)
2
1
0
-1
-2
C
340 downregulated genes
268 upregulated genes
-Log10(adjusted p-value)
Log2(fold change)
15
10
5
0
-5
0
5
D
Contractility of muscle
CLPP
IGF1R
Function of muscle
PPARGC1A
Mitochondrial Dysfunction
Estrogen Receptor Signaling
VEGFA
IGF1
Function of cardiac muscle
PNPLA2
MEF2C
ESRRA
Esrra
PPARG
Cell viability
Organismal death
Cardiac contractility
Contractility of cardiac muscle
Failure of heart
ADAM17
PPARA
TEAD1
TBX5
KDM5A
DNMT3B

**Supplemental Figure 2. Functional analysis reveals increased lipid remodeling in *Acly*$^{KD}$ (MyH6-Cas9$^{+}$) hearts.**

(A) Principal component analysis of normalized RNAseq values of transcripts in control and *Acly*$^{KD}$ heart tissue. n = 3 animals per group.

(B) Unsupervised hierarchical clustering of normalized RNAseq values of transcripts in control and *Acly*$^{KD}$ heart tissue. Heatmaps of z-scores are presented. n=3 animals per genotype.

(C) Volcano plot of gene expression changes in *Acly*$^{KD}$ compared to control mice, based on RNAseq analysis. Log2(Fold Change) is plotted against each gene comparison [–log10(adjusted p-value)]. The numbers of significantly upregulated and downregulated genes are indicated.

(D) Ingenuity pathway enrichment analysis of normalized RNAseq data in control and *Acly*$^{KD}$ heart tissue. n = 3 animals per group. Blue color coding indicates downregulation, while orange color coding indicates upregulation. Arrows indicate regulation and inhibition of interaction genes and pathways.

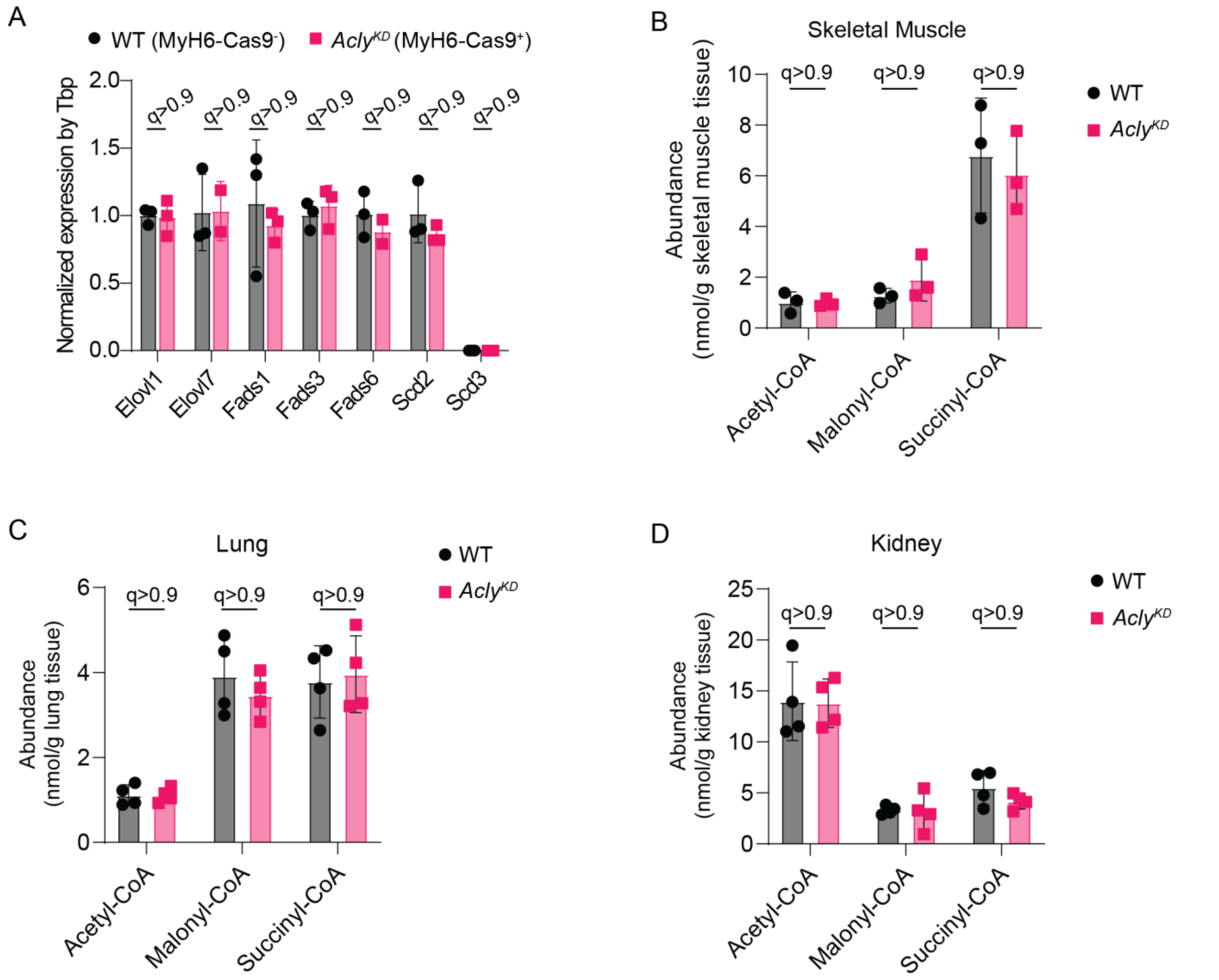


**Supplemental Figure 3. Impact of *Acly$^{KD}$* on lipid metabolism in heart, skeletal muscle, lung, and kidney tissue.**

(A) Gene expression of critical genes for fatty acid synthesis (*Fasn*), desaturation (*Fad2* and *Scd*), and elongation (*Elovl*). Expressions were quantified using RT-qPCR and normalized by *Tbp* expression in each biological sample. n=4-6 male and female mice/group. 2-way ANOVA. Multiple comparison analysis by Sidak.

(B-D) LC-MS/MS-based quantification of acetyl-, malonyl-, and succinyl-CoA in skeletal muscle (M. gastrocnemius) (B), lung (C), and kidney (D) tissue from WT (MyH6-Cas9$^{-}$) or *Acly$^{KD}$* (MyH6-Cas9$^{+}$) mice. n=3-4 male and female mice/group. 2-way ANOVA. Multiple comparison analysis by Sidak.

Numerical q-values are plotted for each comparison. Each data point represents a biological replicate, and error bars represent standard deviation.

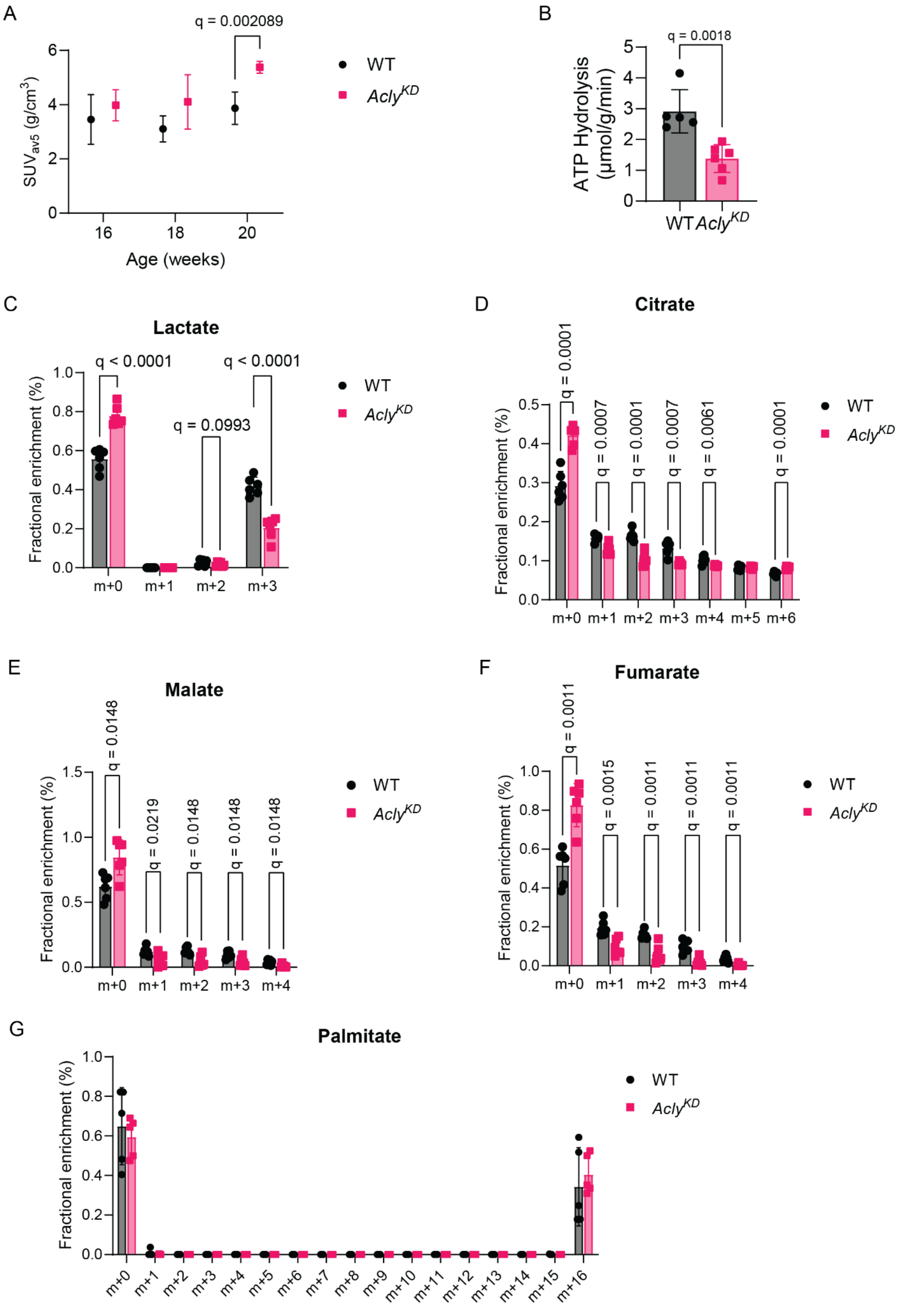
A
q = 0.002089
SUVav5 (g/cm3)
Age (weeks)
WT
AclyKD
B
q = 0.0018
ATP Hydrolysis (µmol/g/min)
WT
AclyKD
C
Lactate
q < 0.0001
q = 0.0993
q < 0.0001
Fractional enrichment (%)
m+0
m+1
m+2
m+3
D
Citrate
q = 0.0001
q = 0.0007
q = 0.0001
q = 0.0007
q = 0.0061
q = 0.0001
m+4
m+5
m+6
E
Malate
q = 0.0148
q = 0.0219
q = 0.0148
q = 0.0148
q = 0.0148
F
Fumarate
q = 0.0011
q = 0.0015
q = 0.0011
q = 0.0011
q = 0.0011
G
Palmitate
m+7
m+8
m+9
m+10
m+11
m+12
m+13
m+14
m+15
m+16

**Supplemental Figure 4. Altered glucose contribution to central carbon metabolism in *Acly*$^{KD}$ hearts.**

(A) Longitudinal [$^{18}$F]-fluorodeoxyglucose (FDG)-positron emission tomography analysis *in vivo* of WT (MyH6-Cas9$^{-}$) or *Acly*$^{KD}$ (MyH6-Cas9$^{+}$) mice at 16, 18, and 20 weeks. Tracer retentions are relative to the heart's weight-normalized injected dose (SUV). The highest five values were used in the average. SUVs were normalized to plasma glucose levels (C). n=7 male and female mice/group. 2-way ANOVA. Multiple comparison analysis by Sidak. FDR<1%.
(B) ATP hydrolysis rate in working heart perfusions from WT (MyH6-Cas9$^{-}$) and *Acly*$^{KD}$ (MyH6-Cas9$^{+}$) mouse hearts at 20 weeks. n = 5-6 male and female mice/group. Statistical significance was calculated by multiple unpaired t-tests followed by multiple comparisons analysis using a false discovery rate < 1% by the two-step method of Benjamini, Krieger, and Yekutieli. Adjusted q-value is blotted for each comparison.
(C-F) Fractional enrichment of the indicated metabolic intermediates from steady-state $^{13}C_6$-glucose perfusions of WT (MyH6-Cas9$^{-}$) and *Acly*$^{KD}$ (MyH6-Cas9$^{+}$) mouse hearts. In all panels, WT data is represented in black, and *Acly*$^{KD}$ data are represented in red. All data are independent measurements from biological replicates. All data are from male and female mice at 20 weeks. n = 6 male and female mice/group. Statistical significance was calculated by multiple unpaired t-tests followed by multiple comparisons analysis using a false discovery rate < 1% by the two-step method of Benjamini, Krieger, and Yekutieli. Adjusted q-value is blotted for each comparison.
(G) Fractional enrichment of $^{13}C_{16}$-palmitate in working heart perfusions of WT (MyH6-Cas9$^{-}$) and *Acly*$^{KD}$ (MyH6-Cas9$^{+}$) mouse hearts at 20 weeks. n = 6 male and female mice/group. Statistical significance was calculated by multiple unpaired t-tests followed by multiple comparisons analysis using a false discovery rate < 1% by the two-step method of Benjamini, Krieger, and Yekutieli. Adjusted q-value is blotted for each comparison.

| Metabolite | Carbons | Formula | m/z |
|---|---|---|---|
| Aspartate | 1,2,3,4 | C4H2NO4 | 418 |
| Citrate | 1,2,3,4,5,6 | C6H4O7 | 459 |
| Fumarate | 1,2,3,4 | C4H2O4 | 287 |
| Glutamate | 1,2,3,4,5 | C5H5NO4 | 432.3 |
| Alanine | 1,2,3 | C3H5NO2 | 160 |
| Glutamine | 1,2,3,4,5 | C5H6N2O3 | 431 |
| Lactate | 1,2,3 | C3H4O3 | 261 |
| Malate | 1,2,3,4 | C4H2O5 | 419 |
| Serine | 1,2,3 | C3H4NO3 | 390 |

**Supplemental Table 2. Metabolite transitions used in metabolic flux analysis.**

| Reaction ID | Reaction | WT | *AclyKD* |
|---|---|---|---|
| **Glycolysis** | | | |
| R1 | Gluc.x -> Gluc | 1 | 1.1265 |
| R2 | Lac -> Pyr | 0 | 0 |
| R3 | Pyr -> Pyr.m | 2 | 2 |
| R4 | Lac -> Lac.x | 0 | 0 |
| R5 | Gluc -> G6P | 1 | 1.1265 |
| R6 | F6P -> FBP | 1 | 1 |
| R7 | PEP -> Pyr | 2 | 2 |
| R8 | G6P -> PG6 | 9E-07 | 0.1265 |
| R9 | PG6 -> Ru5P + CO2 | 9E-07 | 0.1265 |
| R10 | Ru5P -> sink | 9E-07 | 0.1265 |
| R11 | PG3 -> PEP | 2 | 2 |
| R12 | G6P -> F6P | 1 | 1 |
| R13 | DHAP -> GAP | 1 | 1 |
| R14 | GAP -> PG3 | 2 | 2 |
| R15 | FBP -> DHAP + GAP | 1 | 1 |
| **Krebs cycle** | | | |
| R16 | Pyr.m -> AcCoA.m + CO2 | 1.3186 | 1 |
| R17 | Pyr.m + CO2 -> OAA.m | 0.6814 | 1 |
| R18 | OAA.m + AcCoA.m -> Cit.m | 1.3186 | 1 |
| R19 | aKG.m -> Suc.m + CO2 | 0.6372 | 6E-07 |
| R20 | aKG.m -> aKG | 1E-06 | 2.4173 |
| R21 | Cit.m -> aKG.m + CO2 | 0.8113 | 0.1134 |
| R22 | Suc.m -> Fum.m | 0.6372 | 6E-07 |
| R23 | Fum.m -> Mal.m | 0.6372 | 6E-07 |
| R24 | Mal.m -> OAA.m | 1.3361 | 2.314 |
| **Glutamate metabolism** | | | |
| R25 | Gln.x -> Gln | 0.7945 | 0.8 |
| R26 | Gln -> Glu | 0.7945 | 0.8 |
| R27 | aKG -> Glu | 0.6814 | 3.314 |
| R28 | Asp.m + Mal -> Asp + Mal.m | 0.6989 | 2.314 |
| R29 | OAA.m + Glu.m -> Asp.m + aKG.m | 0.6989 | 2.314 |
| R30 | OAA -> Mal | 0.6989 | 2.314 |
| **Citrate metabolism** | | | |
| R31 | Cit.m -> Cit | 1.3803 | 0.8967 |
| R32 | Cit -> aKG + CO2 | 0.6814 | 0.8967 |
| R33 | Cit -> AcCoA + OAA | 0.6989 | 0 |

**Supplemental Table 3. Metabolic flux analysis of WT and *Acly*$^{KD}$ hearts in working mouse heart perfusions using $^{13}C_6$-glucose.**

**Supplemental Figure 5.**

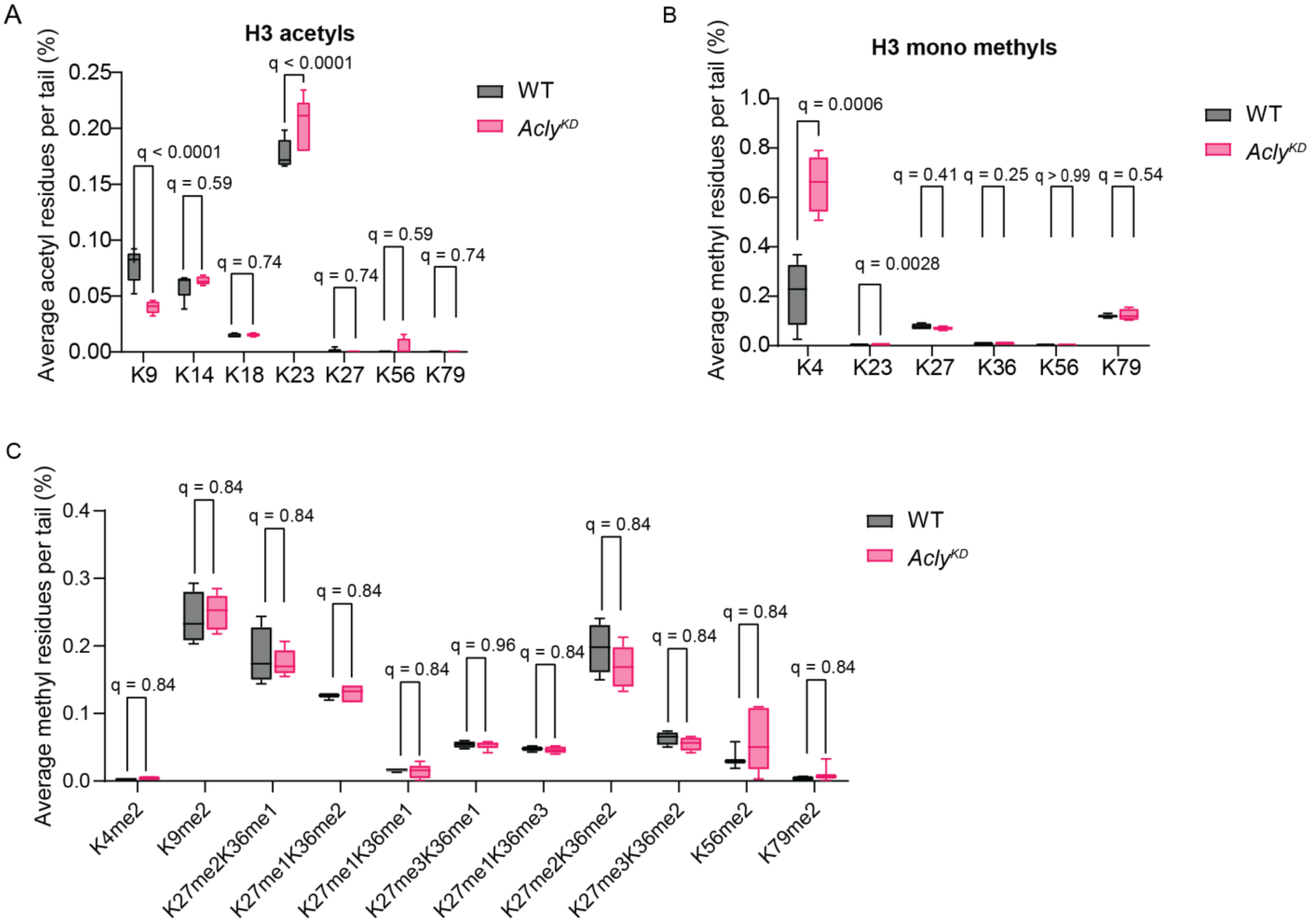


**Supplemental Figure 5. Acly deficiency promotes histone methylation and acetylation changes.**

(A-B) Mass spectrometry-based quantification of the relative abundance of histone 3 acetylation (A) and mono-methylation (B) in heart tissue samples from WT (MyH6-Cas9$^{-}$) or *Acly*$^{KD}$ (MyH6-Cas9$^{+}$) mice at 20 weeks. n=5 male and female mice/group. Statistical significance was calculated by multiple unpaired t-tests followed by multiple comparisons analysis using a false discovery rate <1% by the two-step method of Benjamini, Krieger, and Yekutieli. Adjusted q-value is blotted for each comparison.

(C) Relative abundance of histone 3 di- and tri-methylation residues in heart tissue samples from WT (MyH6-Cas9$^{-}$) or *Acly*$^{KD}$ (MyH6-Cas9$^{+}$) mice at 20 weeks. n=5 male and female mice/group. Statistical significance was calculated by multiple unpaired t-tests followed by multiple comparisons analysis using a false discovery rate < 1% by the two-step method of Benjamini, Krieger, and Yekutieli. Adjusted q-value is blotted for each comparison.

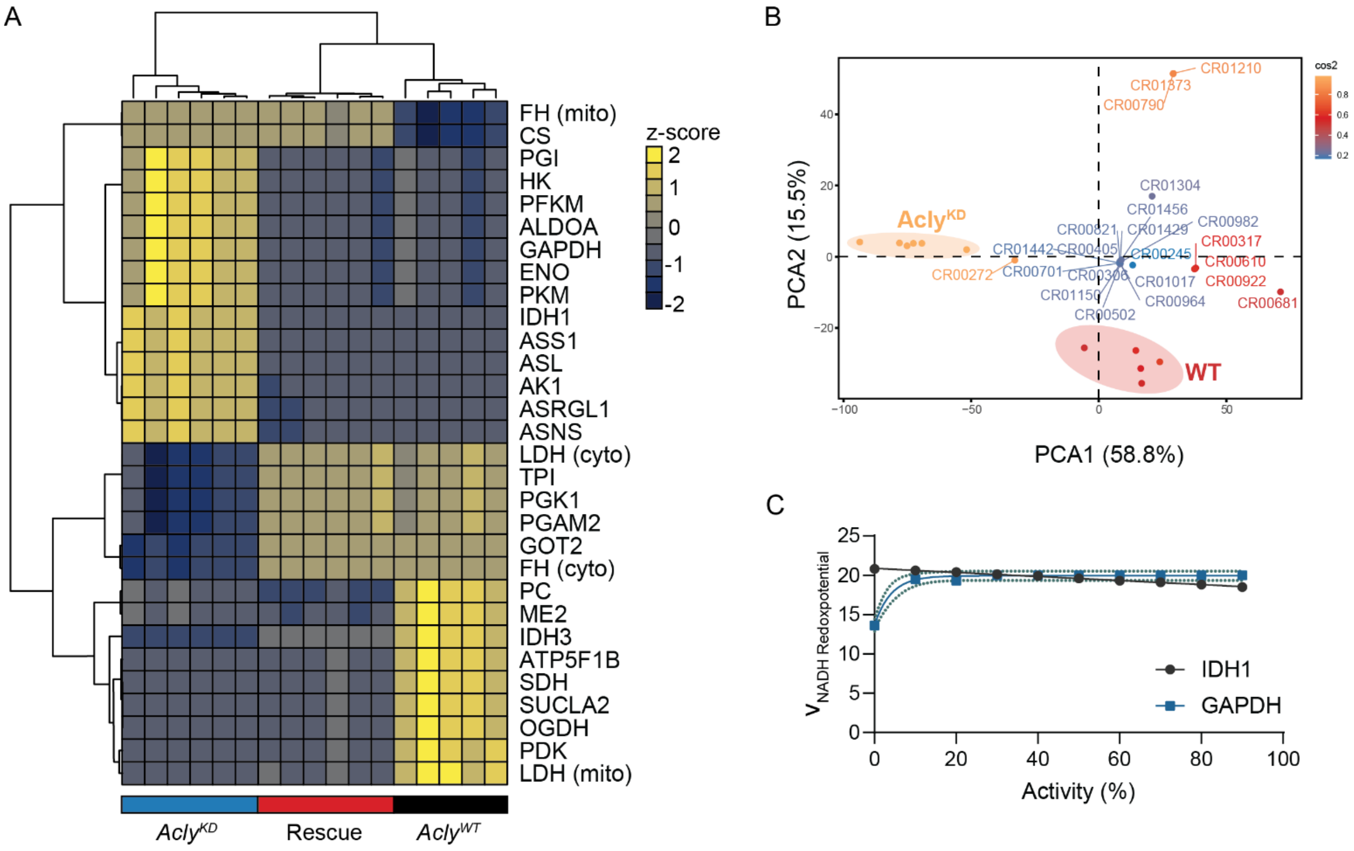


**Supplemental Figure 6. Computational modeling of *Acly*$^{KD}$ and *in silico* rescue simulations.**

(A) Unsupervised hierarchical cluster analysis of significantly altered flux rates in CardioNet simulations for WT (MyH6-Cas9$^{-}$), *Acly*$^{KD}$ (MyH6-Cas9$^{+}$) and *Acly*$^{Rescue}$ simulations.

(B) Principal component analysis of WT (MyH6-Cas9$^{-}$), *Acly*$^{KD}$ (MyH6-Cas9$^{+}$), and *Acly*$^{Rescue}$ simulations.

(D) Relationship between simulated enzyme knockouts for IDH1 and GAPDH, and NADH redox potential in CardioNet simulations.
